# Modulating the properties of DNA-SWCNT sensors using chemically modified DNA

**DOI:** 10.1101/2021.02.20.432105

**Authors:** Alice J. Gillen, Benjamin P. Lambert, Alessandra Antonucci, Daniel Molina-Romero, Ardemis A. Boghossian

**Author notes:** These authors contributed equally to this work (Alice J. Gillen, Benjamin P. Lambert).

## Abstract

Properties of SWCNT-based sensors such as brightness and detection capabilities strongly depend on the characteristics of the wrapping used to suspend the nanotubes. In this study, we explore ways to modify the properties of DNA-SWCNT sensors by using chemically modified DNA sequences, with the aim of creating sensors more suitable for use in *in vivo* and *in vitro* applications. We show that both the fluorescence intensity and sensor reactivity are strongly impacted not only by the chemical modification of the DNA but also by the method of preparation. In the absence of modifications, the sensors prepared using MeOH-assisted surfactant exchange exhibited higher overall fluorescence compared to those prepared by direct sonication. However, we demonstrate that the incorporation of chemical modifications in the DNA sequence could be used to enhance the fluorescence intensity of sonicated samples. We attribute these improvements to both a change in dispersion efficiency as well as to a change in SWCNT chirality distribution.

Furthermore, despite their higher intensities, the response capabilities of sensors prepared by MeOH-assisted surfactant exchange were shown to be significantly reduced compared to their sonicated counterparts. Sonicated sensors exhibited a globally higher turn-on response towards dopamine compared to the exchanged samples, with modified samples retaining their relative intensity enhancement. As the increases in fluorescence intensity were achieved without needing to alter the base sequence of the DNA wrapping or to add any exogenous compounds, these modifications can - in theory - be applied to nearly any DNA sequence to increase the brightness and penetration depths of a variety of DNA-SWCNT sensors without affecting biocompatibility or reducing the near-limitless sequence space available. This makes these sensors an attractive alternative for dopamine sensing *in vitro* and *in vivo* by enabling significantly higher penetration depths and shorter laser exposure times.

## Introduction

The ultimate goal when designing biomedical sensors is to create a technology that can be used for continuous and long-term monitoring applications *in vivo*. In the brain, realizing such technologies could revolutionize health care by improving our understanding and treatment of many different neurological, psychiatric, and cognitive disorders. However, in order to understand both signalling between single neurons and through extended neural networks, it is essential to monitor neural activity over much larger volumes, ideally at high speeds while achieving synaptic resolution^1^. Unfortunately despite significant progress, no single technology or approach has yet been able to accomplish all the analytical requirements for widespread *in vivo* use in humans^2–4^. In addition, for *in vivo* brain imaging, sensors need to be able to transmit through not only tissue and skin, but also trans-cranially. This causes additional attenuation in the fluorescence of optical sensors, creating a need for the removal of overlying brain tissue and for craniotomies to enable sensing in subcortical structures^5^.

An effective strategy for addressing this limitation is to use fluorophores with longer emission wavelengths, such as SWCNTs, where light absorption and scattering by biological tissue is minimised^6–9^. The molecular recognition capabilities of DNA-SWCNTs make them an attractive material for neurotransmitter detection and *in vivo* neural biosensing applications^2,10–16^. However, low photoluminescence quantum yield (QY) values have limited the optical performance of DNA-SWCNTs and constrained their widespread applicability^17–19^ as DNA-SWCNTs typically exhibit much lower fluorescence compared to surfactant-suspended nanotubes^19–23^. As the sensitivity of all nanosensor optical devices is strongly linked to the brightness of the emitting fluorophore^24^, this creates problems for *in vivo* sensing applications. Furthermore, this limits the practical application of the sensors by reducing the penetration depth of the fluorescence emission^25,26^. While there are advantages to using alternative wrappings, like surfactants, to increase the quantum yields of nanotube sensors^23^, ultimately any wrapping replacing DNA sacrifices the increased biocompatibility and vast combinatorial library available for this biopolymer.

Previous attempts to increase DNA-SWCNTs have relied on modifications to the DNA base sequence^25^ or the addition of exogenous compounds. However, many of the exogenous compounds reduce the biocompatibility of the sensors, as the additives (such as reducing agents or nanoparticles) can increase cytotoxicity and diminish sensitivity and/or selectivity of the sensors. Moreover, as the fluorescence properties and molecular recognition capabilities of DNA-SWCNTs have been shown to vary depending on DNA sequence and length ^13,25,27^ there remains a need to identify alternative methods for increasing the fluorescence intensity of DNA-SWCNTs without substituting the wrapping sequence.

Recent work by Yang et al. has demonstrated that the method used to prepare DNA-SWCNT suspensions, direct sonication of methanol (MeOH)-assisted surfactant exchange, can also significantly impact the resulting DNA-SWCNT complex^28^. Disparities in the fluorescence spectra for DNA-SWCNTs produced by the two methods indicated that the replacement process did not always result in the same DNA wrapping configuration compared to using direct sonication. These differences can alter the fluorescence properties of the DNA-nanotube complex but can also impact their recognition capabilities.

In this work, we modulated the properties of (GT)_15_-SWCNT sensors through the incorporation of chemical modifications in the wrapping DNA sequences. The (GT)_15_-SWCNT sensor was selected owing to its important role in neurotransmitter sensing^2,10,12,13^. We examined the impact of these modifications on properties such as dispersion efficiency, fluorescence intensity, and sensor response for SWCNT complexes prepared by both direct sonication and MeOH-assisted surfactant exchange. For sensors prepared by sonication, we observed a notable increase in fluorescence intensity, up to ~79%, when chemically modified DNA was used. Conversely, when using MeOH-assisted surfactant exchange, the incorporation of chemical modifications in the DNA sequence resulted in lower intensities. We further showed that the presence of the modifications did not remove the sensor’s ability to detect dopamine. Moreover, we observed that the response behaviour was strongly dependent on the method of preparation used to construct the DNA-SWCNTs for both modified and non-modified DNA.

These findings demonstrate a new approach for tuning the fluorescence behavior of DNA-SWCNTs for a given sequence and length, without compromising the sensing capabilities of the sensors or requiring the addition of any exogenous compounds.

## Results and Discussion

Modified DNA sequences were chemically synthesized (Microsynth) to contain one of two functional groups: amino or azide (**SI Figure 2, 3**). To investigate the impact of the location and number of modifications on the fluorescence behaviour of the DNA-SWCNTs, modifications were inserted at the 5’ or 3’ end of the DNA sequence, or in the middle of the sequence for (GT)15-SWCNTs. Details on the modifications and positions examined during this study are included in (**Table 1** and **SI Figure 2, 3**).

**Table 1:** A list of functional groups and modification positions tested for improving the fluorescence intensity of (GT)_15_-SWCNTs in this study. Internal modifications for the −NH_2_ functional group were introduced using an amino-dT base (Microsynth) (**SI Figure 3**).

|  | Modification | [Linker] | [Position] | Sequence Name |
| --- | --- | --- | --- | --- |
| (GT)<sub>15</sub> | Unmodified | N/A |  | (GT)<sub>15</sub> |
|  | Amino, - NH<sub>2</sub> | [C<sub>6</sub>] | [5'] | (GT)<sub>15</sub> Amino5' |
|  |  |  | [3'] | (GT)<sub>15</sub> Amino3' |
|  |  |  | [5' + 3'] | (GT)<sub>15</sub> Amino5'3' |
|  |  |  | [5' + Interior] | (GT)<sub>15</sub> Amino5'Int |
|  |  |  | [5' + 3' + Interior] | (GT)<sub>15</sub> Amino5'3'Int |
|  | Azide, -N<sub>3</sub> | [NHS Ester] | [3'] | (GT)<sub>15</sub> Azide5' |
|  |  | [C<sub>6</sub> Amide + C<sub>3</sub> ] | [5'] | (GT)<sub>15</sub> Azide3' |

The absorbance spectra of all (GT)_15_-SWCNTs were used to compare the suspension quality of the solutions that were prepared using direct sonication and MeOH-assisted surfactant exchange (**Figure 1**). All DNA sequences were able to suspend the SWCNTs, as evidenced by the distinct bands in the absorbance spectra and low non-resonant background (**Figure 1 (A)**)^21^. However, variations in the dispersion efficiency were observed depending on both the modification and the preparation method used, as evidenced by large differences in the final SWCNT concentration (post centrifugation, **SI Table 2**). For almost all DNA sequences, higher nanotube yields were obtained for MeOH-assisted surfactant exchange dispersions compared to their sonicated counterparts (**Figure 1** and **SI Table 2**). We also noted that the variation in yields was much greater for DNA-SWCNTs prepared by sonication compared to the surfactant exchange samples. Moreover, while the unmodified (GT)_15_ yielded the most concentrated SWCNT dispersion with the MeOH-assisted surfactant exchange method, the unmodified sequence exhibited one of the lowest yields when prepared by direct sonication, implying that the addition of the functional groups may assist with the exfoliation of the nanotube bundles during the sonication process.

**Figure 1:**
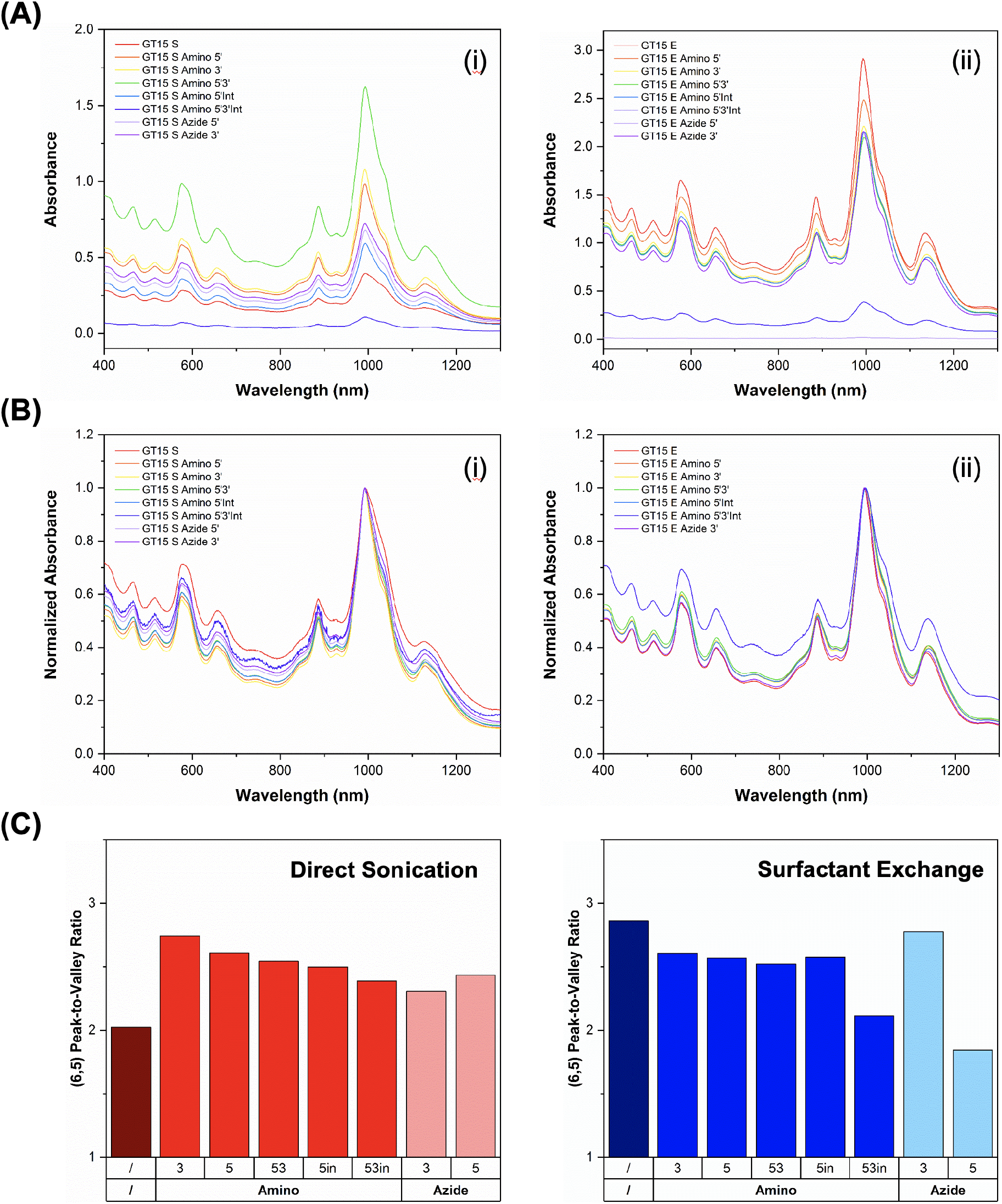
Differences in the suspension quality and yield of solutions examined in this study. **(A)** Absorbance spectra for all modified and unmodified DNA-SWCNT samples prepared via **(i)** direct sonication and **(ii)** MeOH-assisted surfactant exchange. **(B)** Absorbance spectra normalized to the (6,5) E_11_ maximum for all modified and unmodified DNA-SWCNT samples prepared via **(i)** direct sonication and **(ii)** MeOH-assisted surfactant exchange. **(C)** Peak-to-valley ratio for the (6,5) E_11_ peak for the sonicated and exchanged samples.

For both preparation techniques, the Amino 5’3’Int showed a lower final SWCNT concentration. The reduced yield was, in part, attributed to too many modifications in the DNA sequence, which may increase steric hinderance and impact the suspension capabilities of this DNA sequence. Samples containing three modifications displayed the lowest ratios for any modified-DNA samples prepared by both sonication and surfactant exchange, while the singularly modified amino samples showed improved dispersion compared to both the 5’3’ and 5’Int samples for sonicated complexes. We also obtained a very low concentration for the Azide 5’ exchanged sample across multiple preparations (**SI Figure 5**). The difference in suspension quality of Az5’ and Az3’ modifications highlights the impact of the linker molecules, in addition to the functional group, on the dispersion capabilities of a sequence as similar behaviour was also observed when using modified (AT)_15_ sequences (**SI Figure 6**). Due to the lower suspension quality and yields of the samples prepared using Az5’ and Am5’3’Int modifications, these samples were excluded from subsequent fluorescence studies and comparison (**SI Figure 7**).

As expected, the predominant peak in all spectra was observed at around 994 nm, which was associated with the first sub-band E_11_ exciton transition for the (6,5) chirality nanotube. To further compare the samples, the peak-to-valley ratio for the (6,5) absorption peak, which is indicative of relative amounts of impurities and aggregates in solution, was used as a proxy for examining the quality of the suspensions. While we observed improved ratios for all modified DNA sequences in the sonicated samples, the unmodified (GT)_15_ sequence exhibited the highest ratio for the samples prepared by MeOH-assisted surfactant exchange. This again highlights the differences between the two preparation methods and the resultant DNA-SWCNT complexes. Furthermore, we noted a dependence on modification type, position, and abundance, all of which were more pronounced in the sonicated samples. Most notably, we observed a correlation between increasing number of modifications and decreased peak-to-valley ratio.

In addition to variations in absorbance, we observed differences in the fluorescence behaviour of the samples depending on both the abundance and type of modification and the preparation method. We compared fluorescence spectra obtained under an excitation of 575 nm, which is resonant for the (6,5) chirality, for solutions of equal concentration to further examine these differences. For samples prepared by direct sonication, significant increases of up to 79.6% ± 5.3% (**Table 2, SI Table 3**) were obtained in the (6,5) peak fluorescence intensities for all modified (GT)_15_-SWCNTs compared to unmodified (GT)_15_-SWCNTs (**Figure 2 (A), SI Figure 8 (A), 9**). On the other hand, in general the samples prepared by MeOH-assisted surfactant-exchange showed a lower intensity compared to unmodified (GT)_15_ with the exception of the Azide 3’ sample. Interestingly, unmodified (GT)_15_-SWCNTs prepared using MeOH-assisted surfactant exchange displayed a significantly higher fluorescence intensity (55.4 ± 5.6%) compared to those prepared by direct sonication.

**Figure 2:**
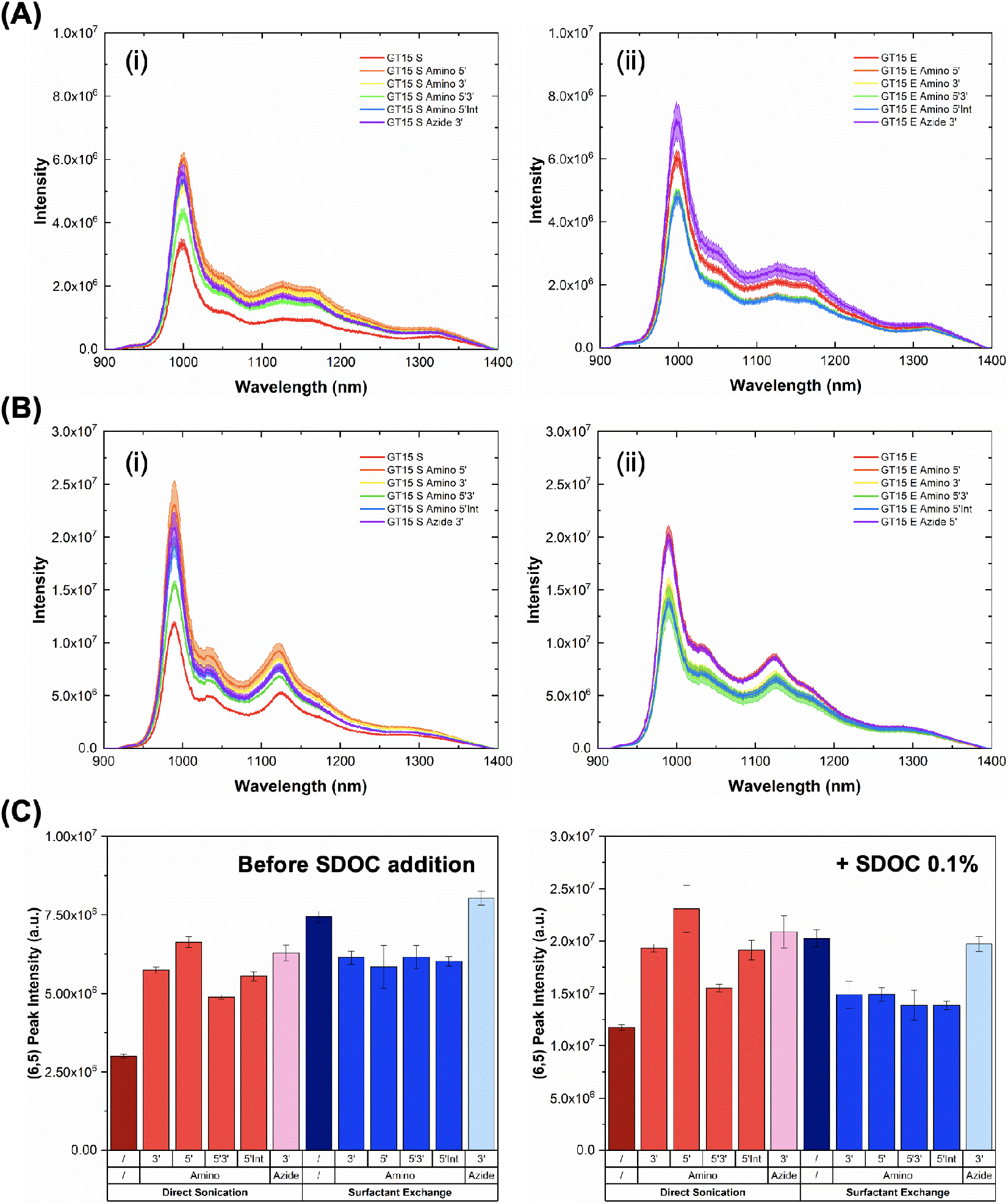
Fluorescence characterisation of SWCNTs suspended with modified and unmodified DNA. **(A)** Fluorescence spectra of the samples suspended via **(i)** direct sonication and **(ii)** MeOH-assisted surfactant exchange (excitation: 575 nm). **(B)** Fluorescence spectra of the samples suspended via **(i)** direct sonication and **(ii)** MeOH-assisted surfactant exchange post replacement by SDOC surfactant (excitation: 575 nm). Graphs include spectra for SWCNTs suspended with unmodified (GT)_15_ (red) and modified (GT)_15_. For all spectra, the central line represents the average spectrum with the shaded regions representing 1*σ* standard deviation (n = 3 technical replicates). All fluorescence spectra were normalized to concentration measured at 739 nm in the 384-well plate immediately prior to measurement to account for any minor variations. **(C)** Comparison of the absolute peak intensity of the (6,5) chirality (excitation: 575 nm) for modified and unmodified (GT)_15_-SWCNTs before and after addition of SDOC (final SDOC concentration: 0.1%). Error bars represent 1*σ* standard deviation (n = 3 technical replicates).

**Table 2:**
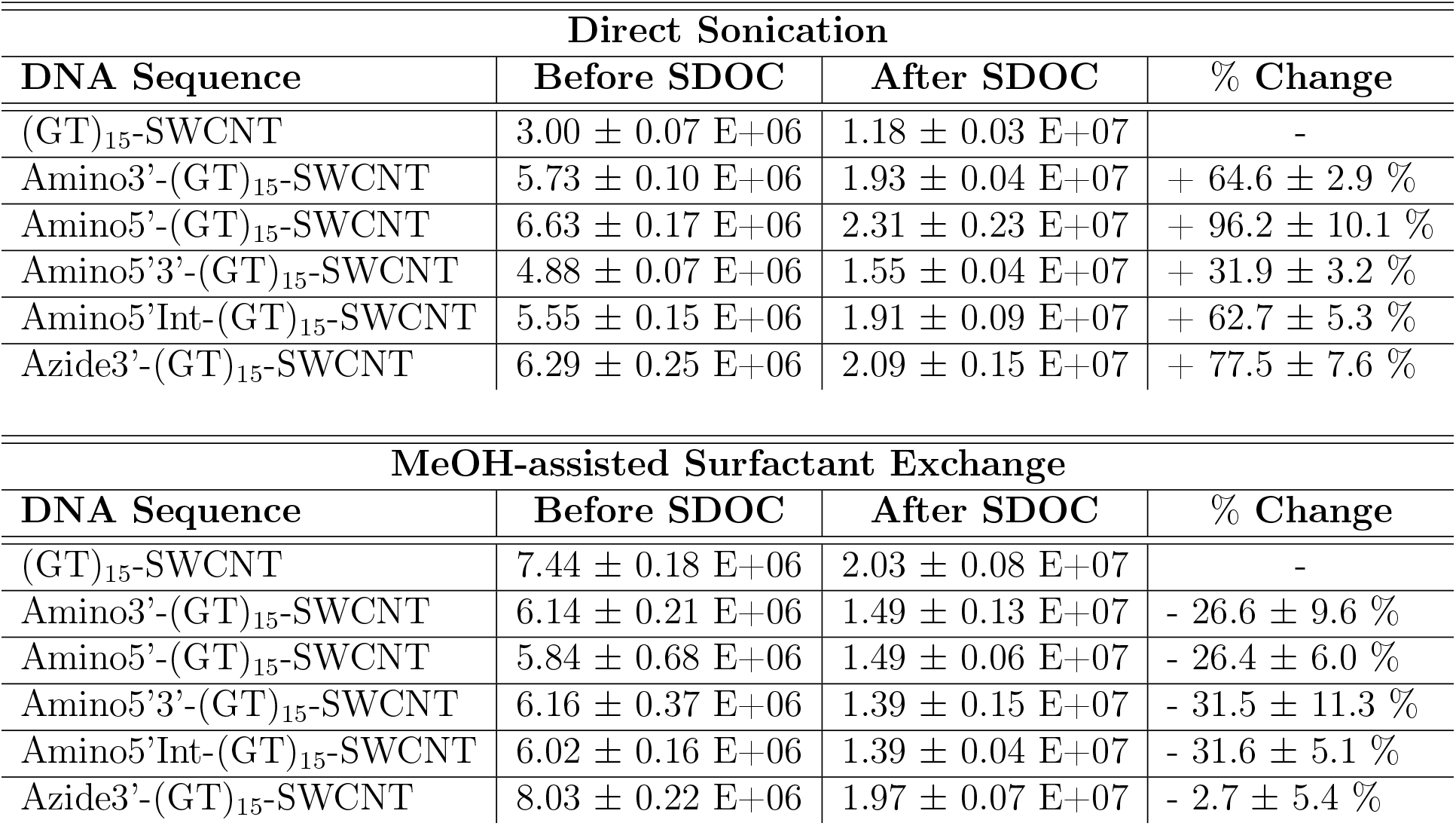
Comparison of the fluorescence intensity of modified and unmodified SWCNTs before and after SDOC addition. Absolute fluorescence intensity of the (6,5) chirality peak for modified and unmodified (GT)_15_-SWCNTs before and after addition of SDOC. Relative intensity changes versus the respective unmodified (GT)_15_-SWCNT complex solution post SDOC addition are also included (% Change).

| Direct Sonication |  |  |  |
| --- | --- | --- | --- |
| DNA Sequence | Before SDOC | After SDOC | % Change |
| (GT)<sub>15</sub>-SWCNT | 3.00 ± 0.07 E+06 | 1.18 ± 0.03 E+07 | - |
| Amino3'-(GT)<sub>15</sub>-SWCNT | 5.73 ± 0.10 E+06 | 1.93 ± 0.04 E+07 | + 64.6 ± 2.9 % |
| Amino5'-(GT)<sub>15</sub>-SWCNT | 6.63 ± 0.17 E+06 | 2.31 ± 0.23 E+07 | + 96.2 ± 10.1 % |
| Amino5'3'-(GT)<sub>15</sub>-SWCNT | 4.88 ± 0.07 E+06 | 1.55 ± 0.04 E+07 | + 31.9 ± 3.2 % |
| Amino5'Int-(GT)<sub>15</sub>-SWCNT | 5.55 ± 0.15 E+06 | 1.91 ± 0.09 E+07 | + 62.7 ± 5.3 % |
| Azide3'-(GT)<sub>15</sub>-SWCNT | 6.29 ± 0.25 E+06 | 2.09 ± 0.15 E+07 | + 77.5 ± 7.6 % |
| MeOH-assisted Surfactant Exchange |  |  |  |
| DNA Sequence | Before SDOC | After SDOC | % Change |
| (GT)<sub>15</sub>-SWCNT | 7.44 ± 0.18 E+06 | 2.03 ± 0.08 E+07 | - |
| Amino3'-(GT)<sub>15</sub>-SWCNT | 6.14 ± 0.21 E+06 | 1.49 ± 0.13 E+07 | - 26.6 ± 9.6 % |
| Amino5'-(GT)<sub>15</sub>-SWCNT | 5.84 ± 0.68 E+06 | 1.49 ± 0.06 E+07 | - 26.4 ± 6.0 % |
| Amino5'3'-(GT)<sub>15</sub>-SWCNT | 6.16 ± 0.37 E+06 | 1.39 ± 0.15 E+07 | - 31.5 ± 11.3 % |
| Amino5'Int-(GT)<sub>15</sub>-SWCNT | 6.02 ± 0.16 E+06 | 1.39 ± 0.04 E+07 | - 31.6 ± 5.1 % |
| Azide3'-(GT)<sub>15</sub>-SWCNT | 8.03 ± 0.22 E+06 | 1.97 ± 0.07 E+07 | - 2.7 ± 5.4 % |

To determine whether the increased fluorescence intensities were due to an increase in quantum yield (QY), we compared the fluorescence intensities of the solutions before and after the addition of excess sodium deoxycholate (SDOC), which shows preferential binding to SWCNTs compared to DNA^29^. After SDOC addition, the fluorescence intensities of all samples increased with a blue-shift in the emission peak, indicative of a total replacement of the DNA wrappings. At the same concentration, SWCNT complexes with equal QY (as a result of SDOC replacement) should exhibit the same fluorescence intensities, evidenced by overlapping spectra^25^. As the peak intensities retained the same trends before and after SDOC addition, we can determine that the fluorescence intensities observed for the modified samples are not exclusively a QY effect (**Figure 2 (C)**).

The retained changes in intensity post SDOC addition are indicative of a change in the SWCNT chirality distribution. Coupled with the variations observed in dispersion quality from the absorbance spectra, we hypothesize that the presence of chemical modifications on the wrapping DNA sequence impacts the dispersion of SWCNTs resulting in changes in the composition and quality of the suspension, which in turn give rise to the differences in fluorescence behaviour. Additionally, despite being prepared with the same batch of surfactant-wrapped nanotubes, samples prepared via MeOH-assisted surfactant exchange also exhibited differences post SDOC-replacement, further supporting the observation that the modified wrappings display different chirality affinities.

Nevertheless, the increase we obtained for modified (GT)_15_-SWCNTs prepared via sonication enables much greater penetration depths compared to their non-modified counterpart. However, as recent studies have shown that DNA sequences can adopt different equilibrium structures depending on their preparation methods^28^, we sought to investigate whether there were differences in the maximal turn-on response following the addition of dopamine for non-modified (GT)_15_-SWCNTs prepared via sonication and surfactant exchange (**Figure 3**). Consistent with previous observations^2,10,23,30^, the addition of dopamine increased the fluorescence intensity of all samples. We define the dopamine response as (I_*f*_-I_0_)/I_0_, where If is the final intensity post-dopamine and I_0_ is the initial intensity. Interestingly, while the initial fluorescence intensity of the (6,5) peak was higher for (GT)_15_-SWCNTs prepared via MeOH-assisted surfactant exchange, we observed a reduction of more than 50% in the turn-on response of the sensor following the addition of dopamine (reduced from 69% to 30%) (**SI Table 4**).

**Figure 3:**
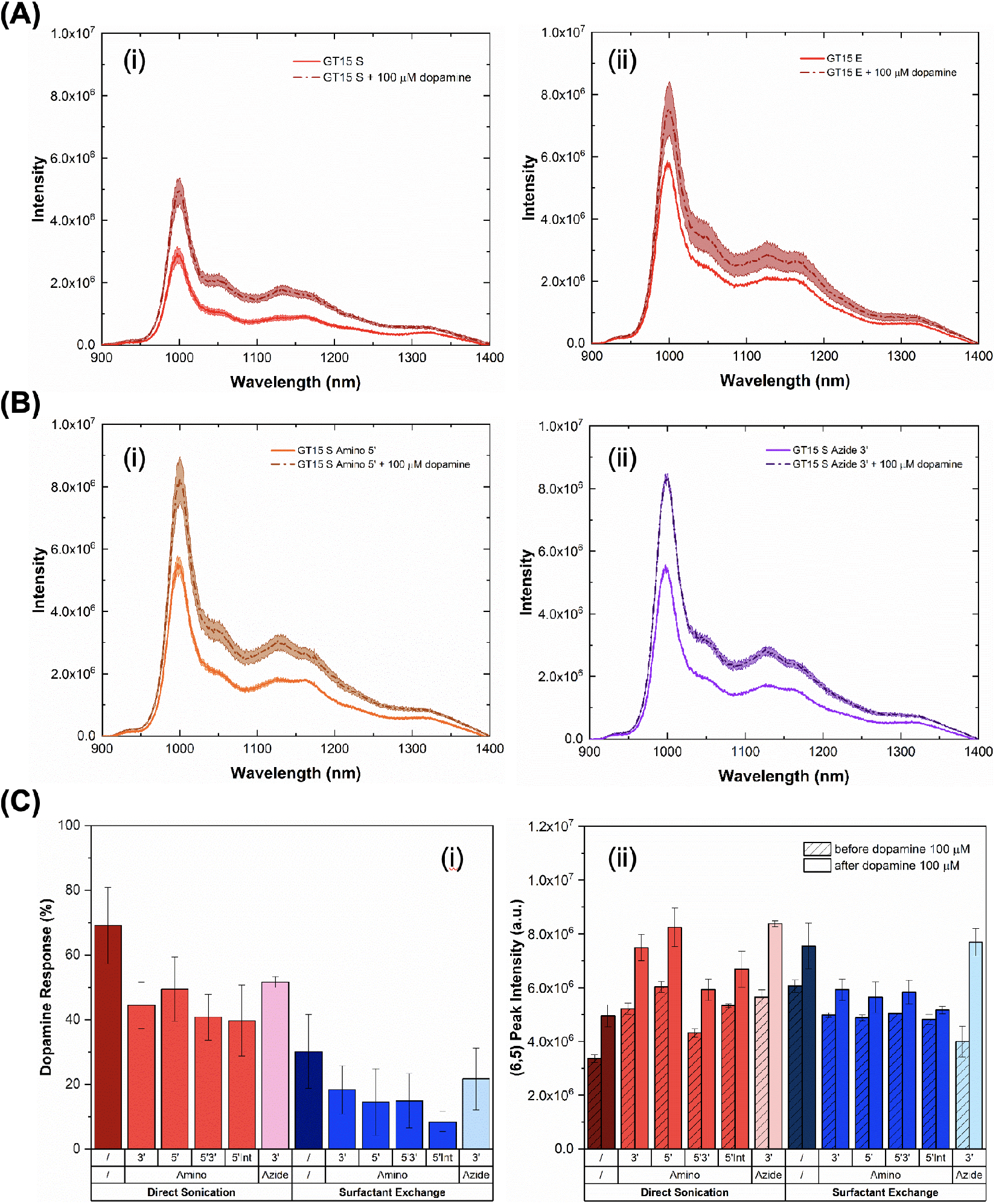
Dopamine response of modified and unmodified (GT) 15-SWCNTs. **(A)** Fluorescence spectra of (GT)_15_-SWCNTs prepared via **(i)** direct sonication and **(ii)** MeOH-assisted surfactant exchange before (solid line) and after (dashed line) addition of dopamine (final concentration: 100 *μ*M, excitation: 575 nm). **(B)** Representative fluorescence spectra of **(i)** Amino 5’- and **(ii)** Azide 3’-modified (GT)_15_-SWCNTs prepared by direct sonication before (solid line) and after (dashed line) addition of dopamine (final concentration: 100 *μ*M, excitation: 575 nm). For all spectra, the central line represents the average spectrum with the shaded regions representing 1*σ* standard deviation (n = 3 technical replicates). **(C)** Fluorescence intensity response following the addition of dopamine. **(i)** Relative intensity response (in %) for all samples. **(ii)** Absolute (6,5) peak intensities before (dashed colour) and after (solid colour) dopamine addition for all samples (final concentration 100 μM, excitation: 575 nm). Error bars represent 1*σ* standard deviation (n = 3 technical replicates).

Moreover, we observed a significantly lower dopamine response for all sensors prepared via surfactant exchange compared to those prepared by direct sonication (**Figure 3 (C), SI Figure 20, 21**). Indeed, all sonicated samples exhibited dopamine responses of at least 2.4 × that of their surfactant-exchanged counterpart. The decrease in dopamine response for these SWCNT suspensions was attributed to the different wrapping configurations of the DNA on the surface of the nanotube^28,31^. Supporting this hypothesis, all nanotube suspensions prepared using MeOH-assisted surfactant exchange also exhibited lower reactivity towards the addition of reducing agents such as dithiothreitol (DTT) (**SI Figure 24 – 28**). These observations suggests that, independent of any chemical modifications, each preparation method resulted in a fundamentally different DNA configuration on the surface of the nanotube, in agreement with recent observations by Yang et al.^28^.

While we did observe a decrease in the turn-on response, the fluorescence enhancement of the modified (GT)_15_-SWCNTs relative to the unmodified sequence was retained post dopamine addition (**Figure 3 (C) (ii), SI Figure 20**). As shown in **SI Table 4**, the absolute intensity increase following dopamine addition (If-I_0_) to the sonicated samples was comparable to the magnitude obtained using unmodified (GT)_15_-SWCNTs and even increased for certain modifications. Moreover, given that both the baseline fluorescence and fluorescence post dopamine addition was higher for modified (GT)_15_-SWCNTs, these sensors could enable *in vivo* sensing at significantly higher penetration depths than unmodified (GT)_15_-SWCNTs (**SI Table 5**)^25,26^, without comprising on the response capabilities of the complexes.

## Conclusion

In this work we examined the impact of chemically modified DNA sequences on the dispersion efficiency and quality of the resulting DNA-SWCNT complexes. We demonstrated that the presence of these modifications can modulate the fluorescence behaviour of the resultant SWCNT complexes, both in terms of fluorescence intensity and response capabilities. Furthermore, we showed that the properties of the DNA-SWCNT complexes, independent of the presence of modifications, are strongly dependent on which preparation method is used.

We observed that the fluorescence intensity of unmodified (GT)_15_-SWCNTs was also significantly lower for sonicated samples compared to those prepared via surfactant exchange. While lower intensities were observed for the modified DNA-SWCNTs prepared using the MeOH-assisted surfactant exchange process, the presence of DNA modifications resulted in higher fluorescence intensities for SWCNT complexes produced by direct sonication. Moreover, for the sonicated samples we noted that optimum improvements were obtained for sequences with only one modification, in particular amino 5’ and azide 3’. Additional experiments using surfactant to replace the DNA wrapping suggested that the observed fluorescence changes were due to changes in chirality distribution and suspension quality, rather than exclusively QY effects. However, this enhancement still enabled significant improvements in the brightness of all modified (GT)_15_-SWCNTs prepared via direct sonication compared to their unmodified counterpart.

We subsequently showed that the sensors responded to analytes differently again depending on both the preparation method used and the presence of chemical modifications in the DNA. This was largely attributed to differences in the secondary structures adopted by the DNA on the surface of the nanotube in agreement with recent findings. The sensors prepared by sonication exhibited higher response to analytes compared to the sensors prepared using the MeOH-assisted surfactant exchange method. In the specific case of dopamine detection, while the turn-on response was reduced for modified DNA-SWCNTs, the relative fluorescence intensity enhancement compared to unmodified DNA for the sonicated samples was retained post addition.

In summary, these findings demonstrate a new approach for tuning the fluorescence behaviour of DNA-SWCNTs for a given sequence and length, without losing the sensing capabilities of the sensors or requiring the addition of any exogenous compounds. Furthermore, the use of chemical modifications can be used to modulate fluorescence intensity while retaining the sensors strong response to analytes of interest. The differences in sensor behaviour observed depending on the preparation method highlights the importance of carefully selecting the preparation method used in order to maximise sensor performance for a desired application. Moreover, we demonstrate that by incorporating chemically modified DNA we can achieve the high fluorescence intensity typically observed for surfactant exchange preparations while retaining the strong response behaviour to target analytes exhibited by sonicated dispersions. Although the exact mechanism for the increase in intensity remains an on-going area of study, we attribute it in part to differences in the final composition of the suspensions in terms of chirality distribution and dispersion quality. We believe that this knowledge can be used as a generic methodology to artificially engineer DNA-SWCNTs for various sensing applications by creating sensors with enhanced fluorescence that enable significantly increased imaging depths *in vivo*.

## Acknowledgement

The authors are thankful for funding support from the Swiss National Science Foundation (SNSF) Assistant Professor (AP) Energy Grant.

This document was prepared as an account of work sponsored by an agency of the United States government. Neither the United States government nor Lawrence Livermore National Security, LLC, nor any of their employees makes any warranty, expressed or implied, or assumes any legal liability or responsibility for the accuracy, completeness, or usefulness of any information, apparatus, product, or process disclosed, or represents that its use would not infringe privately owned rights. Reference herein to any specific commercial product, process, or service by trade name, trademark, manufacturer, or otherwise does not necessarily constitute or imply its endorsement, recommendation, or favoring by the United States government or Lawrence Livermore National Security, LLC. The views and opinions of authors expressed herein do not necessarily state or reflect those of the United States government or Lawrence Livermore National Security, LLC, and shall not be used for advertising or product endorsement purposes.

## Materials and Methods

### Materials

All DNA oligomers were purchased from Microsynth. Chemicals were purchased from Sigma-Aldrich, unless otherwise specified. Purified CoMoCAT SWCNTs were purchased from CHASM Technologies, Inc. (SG65i, batch LOT No. SG65i-L59).

### Preparation of SWCNT Solutions

Suspensions of purified CoMoCAT (Sigma Aldrich SWeNT SG65i, batch MKBN5945V) SWCNTs were prepared using two different methods (1) direct sonication and (2) MeOH-assisted surfactant exchange. All DNA-SWCNT solutions were stored at 4°C between measurements in order to mitigate aggregation of the SWCNTs^32^. Three ssDNA sequences were used in this study, (GT)_15_, (AT)_15_, and N_30_ both unmodified and with various modifications (see Table 1 and SI Table 1 for more information on the modifications used). N_30_ is a random mixture of 30-mer DNA sequences where the exact sequences are not known (1.15 x 10^1^8 unique DNA sequences).

### Direct sonication

1 mg of CoMoCAT SWCNTs was added to a 1 mL solution of ssDNA (100 *μ*M in DI water, Microsynth) and sonicated (140mm, Q700, Qsonica) for 90 min (power = 40 W) in an ice-bath. This was followed by a 4 h centrifugation step (Eppendorf Centrifuge 5424R) at 21,130 × g and 4°C to remove SWCNT aggregates. The supernatant of the suspensions (~ 80% of the solution) was extracted and subsequently washed as detailed below to remove any impurities and unbound DNA from the solution (SI Figure 36).

### MeOH-assisted surfactant exchange

Sodium cholate (SC)-suspended SWCNTs were used for the modified surfactant exchange protocol as previously described^23^. Briefly, 25 mg of CoMoCAT SWCNTs were added to 25 mL of 2% (w/v) SC solution. The mixture was homogenized for 20 min at 5,000 rpm (Polytron PT 1300 D, Kinematica) and subsequently sonicated using probe-tip sonication (1/4 in. tip, Q700 Sonicator, Qsonica) for 1 h (10%amplitude) in an ice bath. The resulting solution was centrifuged at 164,000 × g (30,000 rpm) for 4 h at 20°C (Optima XPN-80 Ultracentrifuge, Beckman) to remove any remaining nanotube aggregates.

To perform the MeOH-assisted surfactant exchange, 400 *μ*L of SC-SWCNTs were mixed with 400 *μ*L of ssDNA solution (75 *μ*M in DI water). The same SC-SWCNT stock was used for all suspensions to ensure a similar starting distribution of nanotube chiralities and lengths for all samples. DNA concentrations were measured and adjusted based on absorbance measurements (Nanodrop 2000, Thermo Scientific). 1.2 mL of methanol (VWR Chemicals) was added to the DNA and SC-SWCNT mixture to obtain a final solvent percentage of 60% (v/v) and a final DNA concentration of 15 *μ*M (SI Figure 37). The solution was vortexed briefly to mix and subsequently incubated for 2 h at room temperature. Following the incubation, all MeOH, displaced surfactant, and unbound DNA was removed by rinsing the solutions according to the procedure detailed below.

### Amicon rinsing for DNA-SWCNT purification

In order to remove impurities (such as remaining catalyst particles, surfactant, MeOH) and unbound DNA from the DNA-SWCNT suspensions, all solutions were purified using Amicon centrifugal ultra-filtration devices (Amicon Ultra-2, Sigma Aldrich, 100 kDa membrane, Merck). Prior to use, the filtration devices were rinsed two times with DI water in accordance with the manufacturer’s recommendations. The solution of DNA-SWCNTs was subsequently added to the filtration device and rinsed eight times with 1 mL aliquots of DI water (devices were centrifuged at 3,000 × g 4°C for 2 min for each rinsing step). The rinsed suspension was collected from the filtration device and centrifuged for a minimum of 1 h at 21,130 × g and 4°C to remove any additional SWCNT aggregates that may have formed during the rinsing process. The supernatant solution was extracted and subsequently characterised using a UV-Vis-NIR scanning spectrometer (Shimadzu 3600 Plus) with a quartz cuvette (Suprasil quartz, path length 3 mm, Hellma).

### Absorption spectroscopy

Absorbance spectra were acquired for all samples using a UV-Vis-NIR scanning spectrometer (UV-3600 Plus, Shimadzu) with a quartz cuvette (Suprasil quartz, path length 3 mm, Hellma). The concentrations for all samples were calculated using an extinction coefficient ɛ_739*nm*_ = 0.0253 L mg^-1^ cm^-13 3^. All nanotube suspensions were diluted to 33 mg/L, corresponding to Abs_739*nm*_ = 0.253, unless otherwise specified.

SDOC replacement samples were prepared by mixing 90 *μ*L of SWCNT solution (~ 33 mg/L) with 10 *μ*L of SDOC 1% (w/v). Spectra for the samples were collected prior to addition and following a period of 10 min incubation post-addition.

### Fluorescence spectroscopy measurements

Fluorescence emission spectra were acquired using a custom-built optical set-up with an inverted Nikon Eclipse Ti-E microscope (Nikon AG Instruments), as described previously^34^. Briefly, samples were excited using a pulsed super-continuum laser coupled with a tuneable band-pass filter unit (SuperK Extreme EXR-15 and Super K VARIA, NKT Photonics). The fluorescence signal was collected using an IsoPlane SCT-320 spectrometer (Princeton Instruments) coupled to an InGaAs NIR camera (NIRvana 640 ST, Princeton Instruments). Measurements were recorded with LightField (Princeton Instruments) in combination with a custom-built LabView (National Instruments) software for automation purposes. An exposure time of 5 s and laser excitation with band width of 10 nm and relative power of 100 %was used for all measurements, unless stated otherwise. Fluorescence emission spectra were collected at wavelengths between 900 nm and 1400 nm using a dispersive grating of 75 lines mm^-1^. All experiments were performed in 384-well plates (Clear Flat-Bottom Immuno Nonsterile 384-Well Plates, MaxiSorp, Life Technologies) which were sealed (Empore Sealing Tape Pad, 3M) prior to each fluorescence measurement to prevent evaporation.

### Photoluminescence measurements

50 *μ*L aliquots of DNA-SWCNT solutions (~ 33 mg/L) were added to a 384-well plate and photoluminescence excitation (PLE) maps were acquired between 500 nm and 800 nm using a 5 nm step and 5 s exposure time. Additional samples were used to acquire on-resonance fluorescence emission spectra for the (6,5) chirality (at 575 ±5 nm) and (7,5) chirality (at 660 ±5 nm). Absorbance values at 575, 632, 660, 739, and 808 nm were collected immediately prior to measurement using a Varioskan LUX microplate reader to ensure that the concentrations of the nanotubes were comparable. All spectra were normalised to the concentration determined at Abs_739*nm*_. Results were analysed using a custom Matlab code (Matlab R2017b, Mathworks) for the PLE maps and custom Python codes for the on-resonance spectra.

### Dopamine detection assay

10 mM solutions of dopamine (dopamine hydrochloride) were freshly prepared in DI water immediately prior to measurement. Fluorescence spectra were initially acquired for all DNA-SWCNT solutions (49.5 *μ*L, ~ 33 mg/L) in a 384-well plate using a laser excitation of 575 ±5 nm. Following the initial measurement, 0.5 *μ*L of dopamine solution (10 mM) was added to the SWCNT suspension. Solutions were mixed by pipetting up and down several times. The suspensions were incubated for 10 min at room temperature in the dark prior to recording the second fluorescence spectrum.

### Surfactant replacement assay

Surfactant replacement of the DNA-SWCNT solutions was performed using 1% (w/v) SDOC. 5 *μ*L of SDOC was mixed with 45 *μ*L aliquots of the DNA-SWCNT solutions (~ 33 mg/L) and incubated at room temperature for 10 min. Fluorescence spectra were acquired before and after SDOC addition using an excitation of 575 ±5 nm.

### pH and DTT assay

Baseline fluorescence spectra were acquired for all DNA-SWCNT solutions (49.5 *μ*L, ~ 33 mg/L) in a 384-well plate using an excitation of 575 *±*5 nm. Following this, 0.5 μL of either NaOH 0.02 M, NaOH 0.1 M, or DTT was added to the SWCNT suspension. Solutions were mixed by pipetting up and down several times. Prior to recording the second spectrum, suspensions were incubated for 10 min at room temperature.

### Confocal Raman microscopy

Samples were prepared for Raman microscopy by drop casting 10 *μ*L of DNA-SWCNT solution (10 mg/L) onto cleaned glass slides (Coverslip 24×55×0.15 mm, Fisher Scientific). Raman spectra were recorded at an excitation wavelength of 532 nm at 100%relative power with 5 s exposure time. The spectrometer was calibrated before measurements using an internal standard. Spectra were collected between 100 – 2750 cm^-1^ using a water-immersion 100×L objective on a confocal spectroscope (inVia Raman Microscope, Renishaw) with a grating of 1,800 lines mm^-1^.

### Gel electrophoresis

Unbound DNA was removed from DNA-SWCNT samples prepared by direct sonication and MeOH-assisted surfactant exchange by rinsing with Amicon centrifugal ultra-filtration devices (Amicon Ultra-2, Sigma Aldrich, 100 kDa membrane, Merck), as detailed above. Samples were diluted to a concentration of 1 mg/L (extinction coefficient ɛ_739*nm*_= 0.0253 L mg^-1^cm ^-1 33^) and DNA was extracted using phenol-chloroform isoamyl (PCI). Equal volumes of DNA-SWCNT suspension and PCI solution (Sigma-Aldrich 25:24:1) were combined and vortexed for 30 s. The solution was centrifuged at 16,000 × g for 5 min (at room temperature). The aqueous top phase was collected and DNA was subsequently precipitated by ethanol precipitation in the presence of sodium acetate and glycogen (Carl Roth, final concentration of 1 g/L)^35^. The DNA pellet was washed with 70% ethanol to remove salts and finally resuspended in 10 *μ*L of DI water.

Extracted DNA was mixed with equal volumes of 2 × formamide loading buffer. The samples were denatured by incubating the mixture at 95°C for 5 min and immediately quenching it on ice before loading the solution onto the gel. The samples were run on a denaturing 15% urea-polyacrylamide gel in 1 × Tris/Borate/EDTA (TBE) buffer at 200 V for 1 h. Three dilutions (1 ×, 0.5 ×, and 0.2 ×) were run for each DNA sequence. SYBR Gold dye (0.2 ×, Thermo Fisher) as used to stain the DNA. Fluorescence was recorded following 30 min of staining on a blue-light gel image (E-Gel, Thermo Fisher). In order to compare the size of the DNA fragments, a ssDNA ladder (10 – 100 nucleotides) was run in the last lane of the gel.

### Supporting Information Available

Data supporting the findings of this study are available within the paper and its Supplementary Information file. All other relevant data and other findings of this study are available from the corresponding author upon reasonable request.

