## Supplementary information for "Modulating the properties of DNA-SWCNT sensors using chemically modified DNA"

### DNA-dimer formation of thiol-modified DNA

Over time, thiol-modified DNA oxidizes, resulting in the formation of disulfide bonds between two thiol groups on the sequences. Polyacrylamide gel electrophoresis (PAGE) analysis of DNA-sequences removed from DNA-SWCNTs post-suspension showed that thiol-modified DNA formed DNA dimers during the suspension process (**SI Figure 1**), resulting in an effective sequence length of 60 bases (compared to the 30 bases for the amino-, azide-, and unmodified sequences). Although reduction steps can be used to break the disulfide bonds and separate oligo dimers, reducing agents (such as dithiothreitol (DTT)) can significantly affect the fluorescence properties of DNA-SWCNTs. As a result, in order to ensure accurate comparisons between the functionalizations, thiol-modified DNA sequences were excluded from this study.

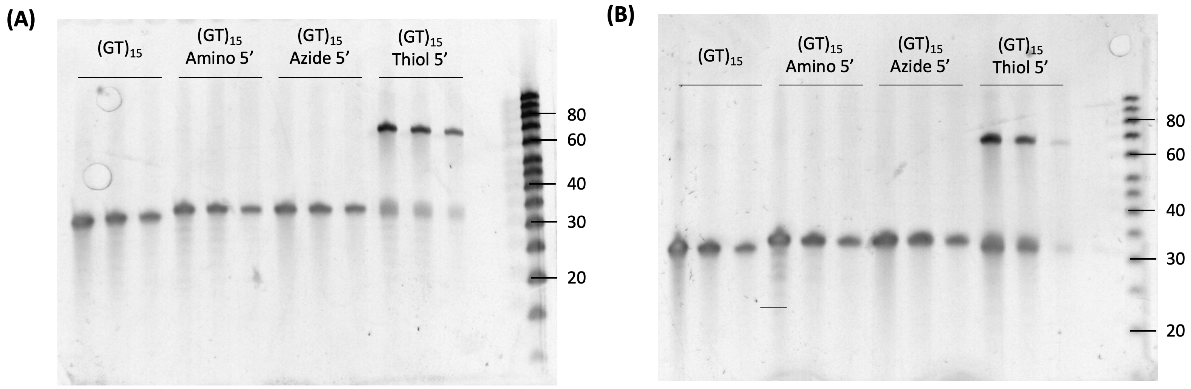


Figure 1: Denaturing 15% polyacrylamide gels of DNA extracted from **(A)** sonicated and **(B)** MeOH assisted surfactant exchanged DNA-SWCNT samples. The DNA samples were extracted by phenol-chloroform isoamyl (PCI) extraction and subsequently precipitated through glycogen-assisted ethanol precipitation. DNA samples were run at three concentrations for each sequence: $1\times$, $0.5\times$, and $0.2\times$ in the first, second, and third lanes, respectively. Both gels were stained using a SYBR Gold dye staining. A band around 30 nucleotides is observed as expected for every sequence. An additional band at $\sim 60$ nucleotides is also present for the thiol-modified DNA sequence, indicating the presence of dimers.

**Table 1:** Additional DNA sequences, functional groups, and modification positions tested for improved fluorescence intensity of nanotube suspensions prepared using the MeOH assisted surfactant exchange protocol.

| | **Functional group** | **[Linker]** | **[Modification Positions]** |
|---|---|---|---|
| $(AT)_{15}$ | Amino ($-NH_2$) | [$C_6$] | [Unmodified] [5'] [3'] [5' + 3'] |
| | Azide ($-N_3$) | [NHS Ester] | [5'] [3'] |
| $(N)_{30}$ | Amino ($-NH_2$) | [$C_6$] | [Unmodified] [5'] [5' + 3'] |
| | Azide ($-N_3$) | [NHS Ester] | [5'] |

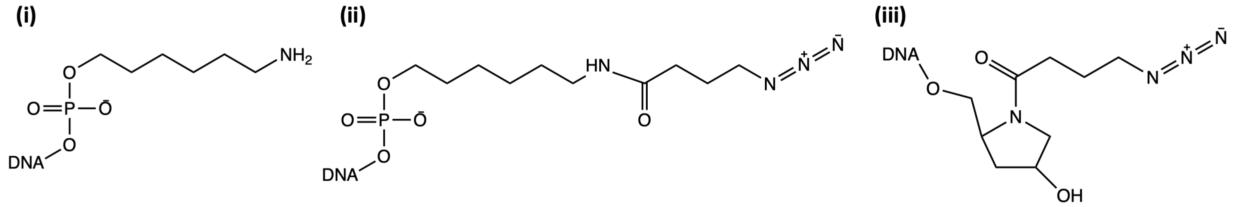


Figure 2: Chemical structures of the functional groups and linker molecules used for terminal DNA modification: **(i)** amino, **(ii)** azide 5' and **(iii)** azide 3'. All chemical structures were drawn in ChemDraw Prime 17.0.

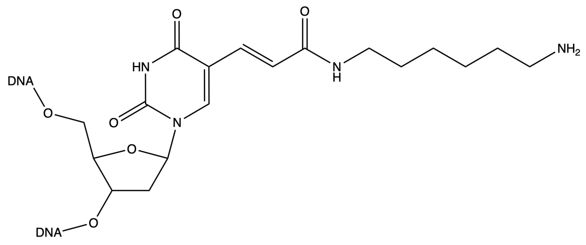


Figure 3: Chemical structure of amino-dT base used for internal modification of DNA sequences with an amino functional group drawn using ChemDraw Prime 17.0.

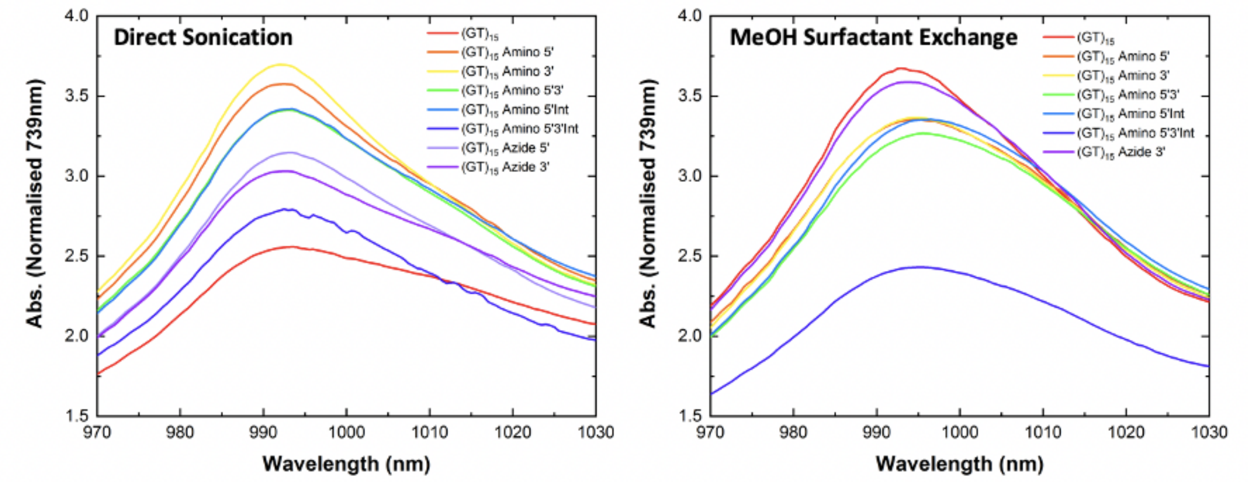


Figure 4: Comparison of the concentration normalized absorbance spectra for the (6,5) chirality $E_{11}$ peak of unmodified and modified $(GT)_{15}$-SWCNTs prepared via (**left**) direct sonication and (**right**) MeOH assisted surfactant exchange. All spectra were normalized to concentration using an extinction coefficient of 0.0253 mg/L at $Abs_{739nm}$.

**Table 2:** A comparison of the concentration yield of SWCNTs suspended using direct sonication and MeOH-assisted surfactant exchange for both modified and unmodified $(GT)_{15}$.

| **Sequence** | **Direct Sonication** (mg/L) | **MeOH-assisted Surfactant Exchange** (mg/L) |
|---|---|---|
| $(GT)_{15}$ | 40.74 | 209.01 |
| $(GT)_{15}$ Amino 5’ | 72.50 | 195.01 |
| $(GT)_{15}$ Amino 3’ | 77.10 | 173.19 |
| $(GT)_{15}$ Amino 5’3’ | 125.50 | 168.91 |
| $(GT)_{15}$ Amino 5’Int | 45.60 | 169.08 |
| $(GT)_{15}$ Amino 5’3’Int | 10.22 | 41.99 |
| $(GT)_{15}$ Azide 5’ | 56.96 | 2.67 |
| $(GT)_{15}$ Azide 3’ | 62.84 | 158.36 |

### Azide-modified $(GT)_{15}$

We observed a significant difference between $(GT)_{15}$ Azide5'-SWCNTs and $(GT)_{15}$ Azide3'-SWCNTs for suspensions prepared using MeOH-assisted surfactant exchange. While $(GT)_{15}$ Azide3' continued to produce well dispersed, high-yield solutions of SWCNTs with peak-to-background ratios similar to unmodified $(GT)_{15}$, $(GT)_{15}$ Azide5' was not able to effectively suspend the nanotubes. The yield of $(GT)_{15}$ Azide5'-SWCNTs was more than sixty times lower in concentration compared to the $(GT)_{15}$ Azide3'-SWCNTs and had the highest FWHM and lowest relative absorbance of all solutions. Subsequent preparations of $(GT)_{15}$ Azide5'-$(GT)_{15}$-SWCNTs using the surfactant exchange protocol resulted in equally low-yields (**SI Figure 5**). Similarly, decreases in the yield and absorbance of the (6,5) peak were observed for nanotube suspensions prepared via surfactant exchange with $(AT)_{15}$ Azide5' but not $(AT)_{15}$ Azide3' (**SI Figure 6**). The difference in suspension quality of Azide5' and Azide3' modifications highlights the impact of the linker molecules as well as the functional group on the dispersion capabilities of a sequence.

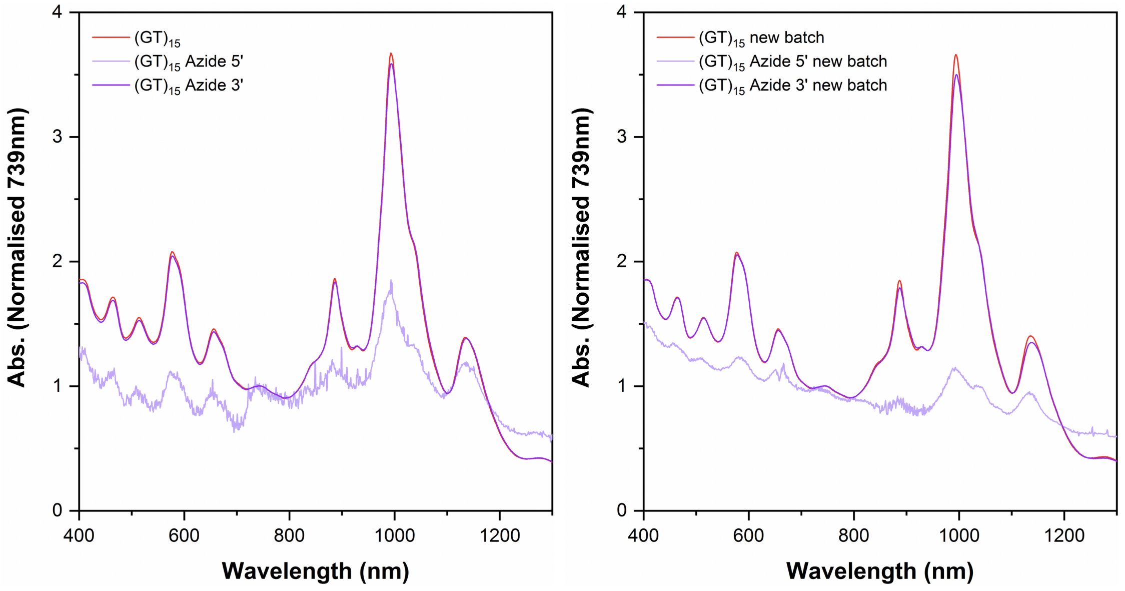


Figure 5: Absorbance spectra for Azide5'- and Azide3'-$(GT)_{15}$-SWCNTs prepared using MeOH assisted surfactant exchange. Two preparations of unmodified, Azide5'-, and Azide3'-$(GT)_{15}$-SWCNTs confirm the much lower dispersion capabilities of Azide5'-$(GT)_{15}$ when the MeOH assisted surfactant exchange protocol is used. All spectra were normalized to concentration using an extinction coefficient of 0.0253 mg/L at $Abs_{739nm}$.

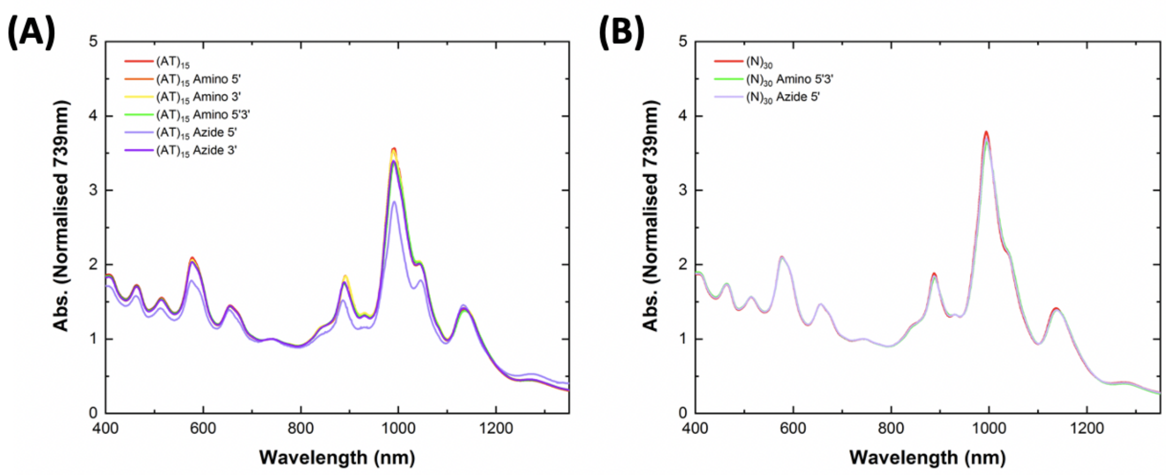


Figure 6: Differences in the suspension quality and chirality distribution of modified and unmodified $(AT)_{15}$- and $(N)_{30}$-SWCNTs prepared using MeOH assisted surfactant exchange. (A) Absorbance spectra for modified and unmodified **(A)** $(AT)_{15}$-SWCNT samples and **(B)** $(N)_{30}$-SWCNT samples. All spectra are normalized to concentration using extinction coefficient of 0.0253 mg/L at $Abs_{739nm}$.

**Table 3:** Fluorescence intensity change of the (6,5) and (7,5) chiralities for modified $(GT)_{15}$-SWCNTs compared to the respective $(GT)_{15}$-SWCNT complex. The integrated fluorescence intensity under the 575 nm excitation is labelled as ALL.

| **DNA Sequence** | **Direct Sonication** | | | **MeOH-assisted Surfactant Exchange** | | |
|---|---|---|---|---|---|---|
| | **(6,5)** | **(7,5)** | **ALL** | **(6,5)** | **(7,5)** | **ALL** |
| $(GT)_{15}$ Amino3' | 55.4 ± 5.8% | 136.6 ± 2.0% | 70.7% | -17.8 ± 4.1% | -30.2 ± 2.4% | -18.5% |
| $(GT)_{15}$ Amino5' | 79.6 ± 5.3% | 132.1 ± 7.2% | 87.2% | -19.6 ± 4.4% | -23.6 ± 2.2% | -18.4% |
| $(GT)_{15}$ Amino5'3' | 28.7 ± 5.5% | 90.6 ± 6.2% | 44.1% | -17.0 ± 3.8% | -27.4 ± 2.0% | -16.7% |
| $(GT)_{15}$ Amino5'Int | 58.7 ± 4.4% | 89.1 ± 1.4% | 62.0% | -20.6 ± 5.3% | -30.6 ± 3.4% | -20.5% |
| $(GT)_{15}$ Azide3' | 67.9 ± 6.4% | 77.3 ± 4.5% | 63.6% | 19.5 ± 8.5% | 27.9 ± 6.9% | 18.4% |

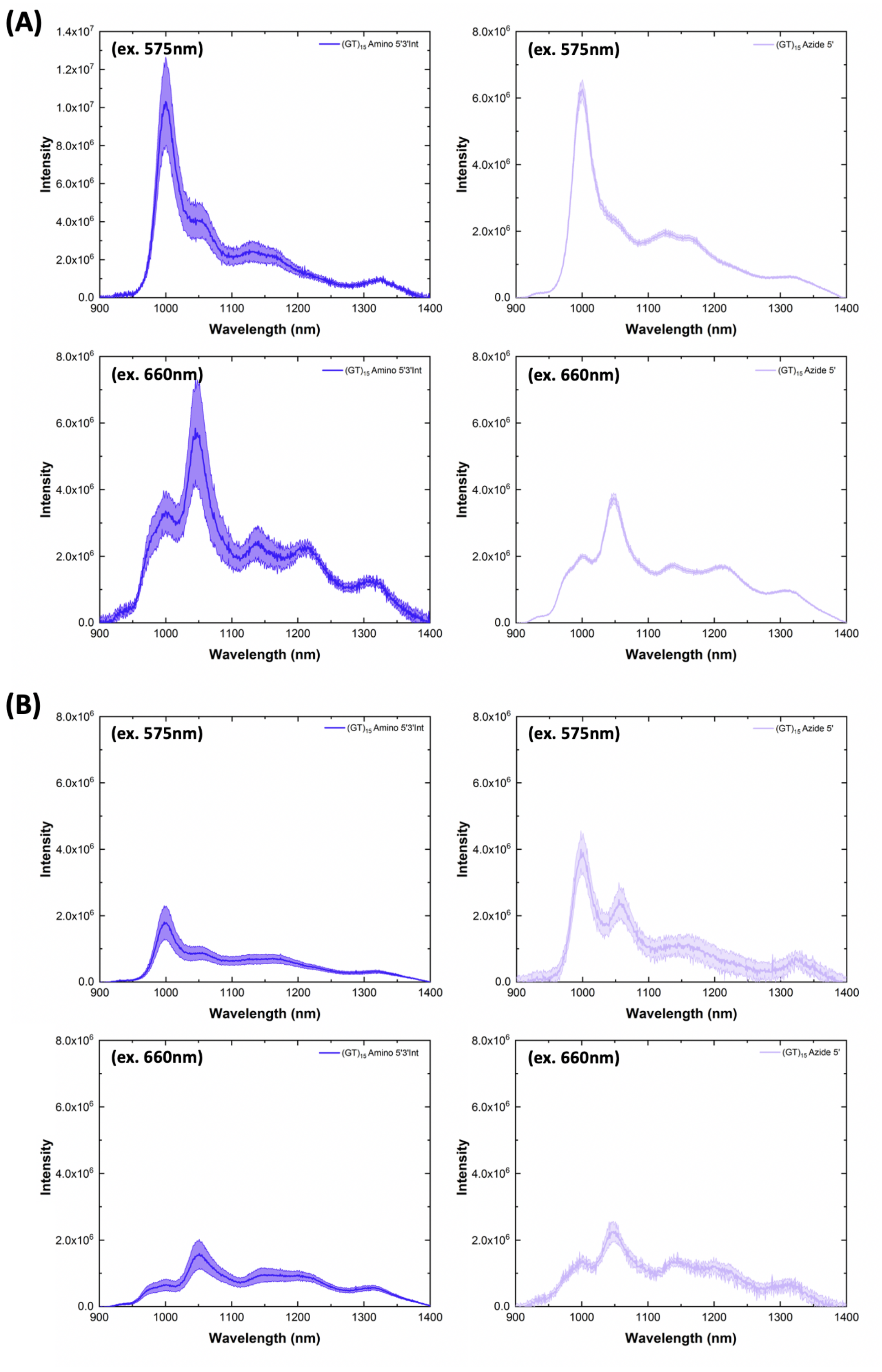


Figure 7: Fluorescence emission spectra for Am5'3'Int-$(GT)_{15}$-SWCNTs (**left**) and Azide5'-$(GT)_{15}$-SWCNTs (**right**) samples prepared by **(A)** direct sonication and **(B)** MeOH assisted surfactant exchange. All fluorescence spectra were normalized to nanotube concentration measured in the 384-well plate immediately prior to measurement to account for any minor variations. For all spectra, the central line represents the average spectrum with the shaded regions representing $1\sigma$ standard deviation (n = 3 technical replicates) (excitation: 575 nm (**top**) and 660 nm (**bottom**)).

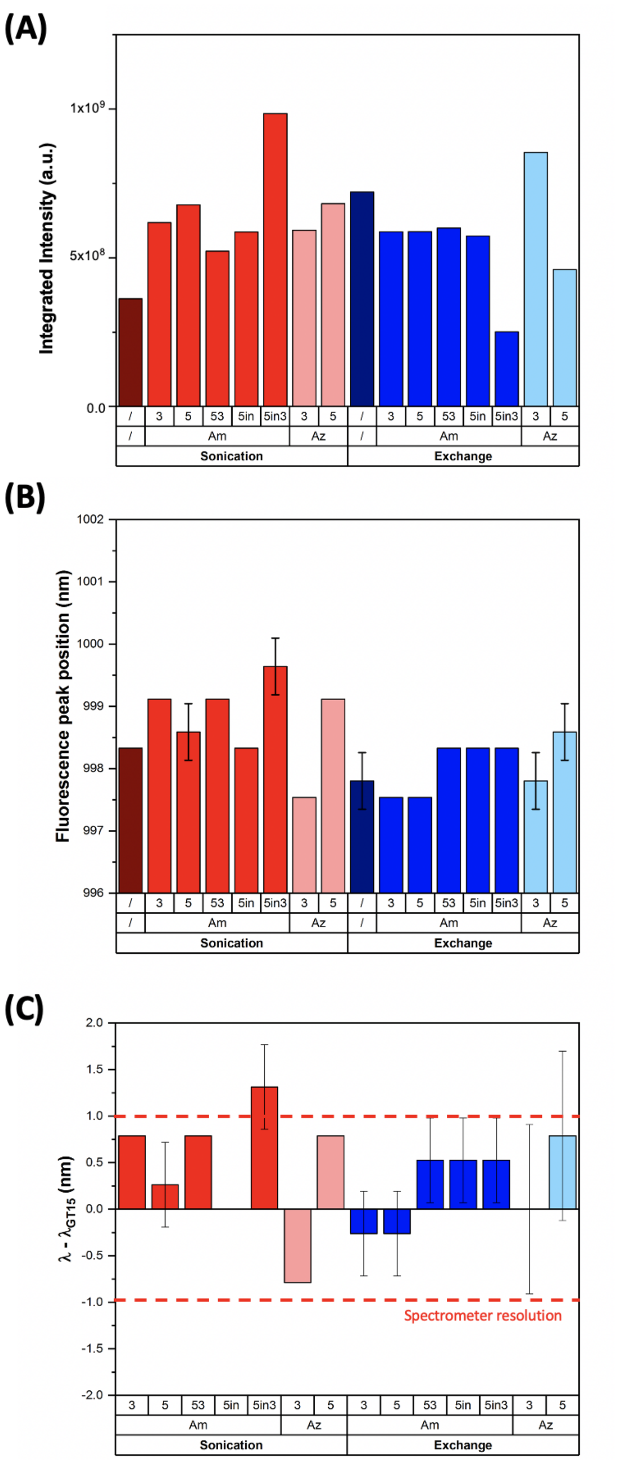


Figure 8: **(A)** Integrated fluorescence intensity under 575 nm excitation of modified and unmodified $(GT)_{15}$-SWCNTs prepared by direct sonication and MeOH assisted surfactant exchange. The integrated intensity was calculated from the additive emissions for wavelengths 900 – 1400 nm. Error bars represent $1\sigma$ standard deviation (n = 3 technical replicates). **(B)** Fluorescence peak positions for the (6,5) chirality (excitation: 575 nm) for all modified and unmodified $(GT)_{15}$-SWCNTs. **(C)** Shift in the peak positions for the (6,5) peak compared to the respective $(GT)_{15}$-SWCNT suspension ($\lambda_{(GT)15}$). Error bars represent $1\sigma$ standard deviation (n = 3 technical replicates). Red dashed lines represent the limit of resolution for the spectrometer.

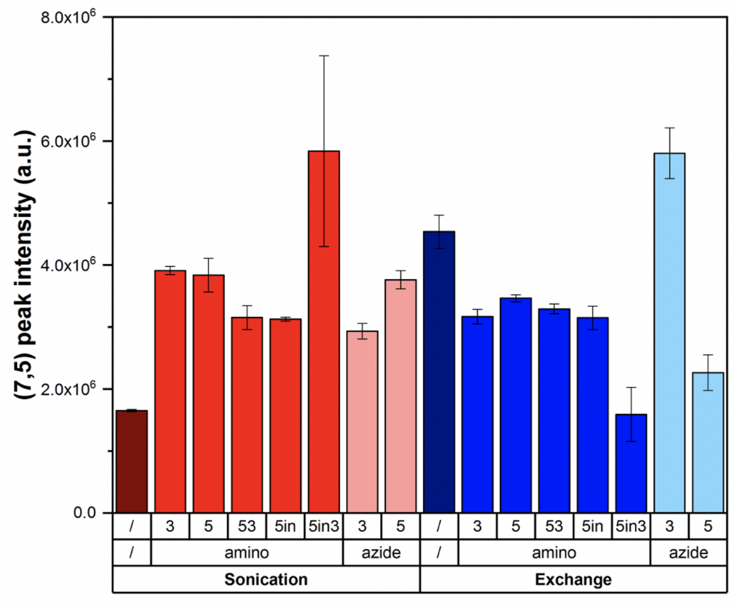


Figure 9: Comparison of the absolute peak intensity of the (7,5) chirality (excitation: 660 nm) for SWCNTs suspended with both modified- and unmodified-$(GT)_{15}$ by direct sonication and MeOH assisted surfactant exchange. Error bars represent $1\sigma$ standard deviation (n = 3 technical replicates).

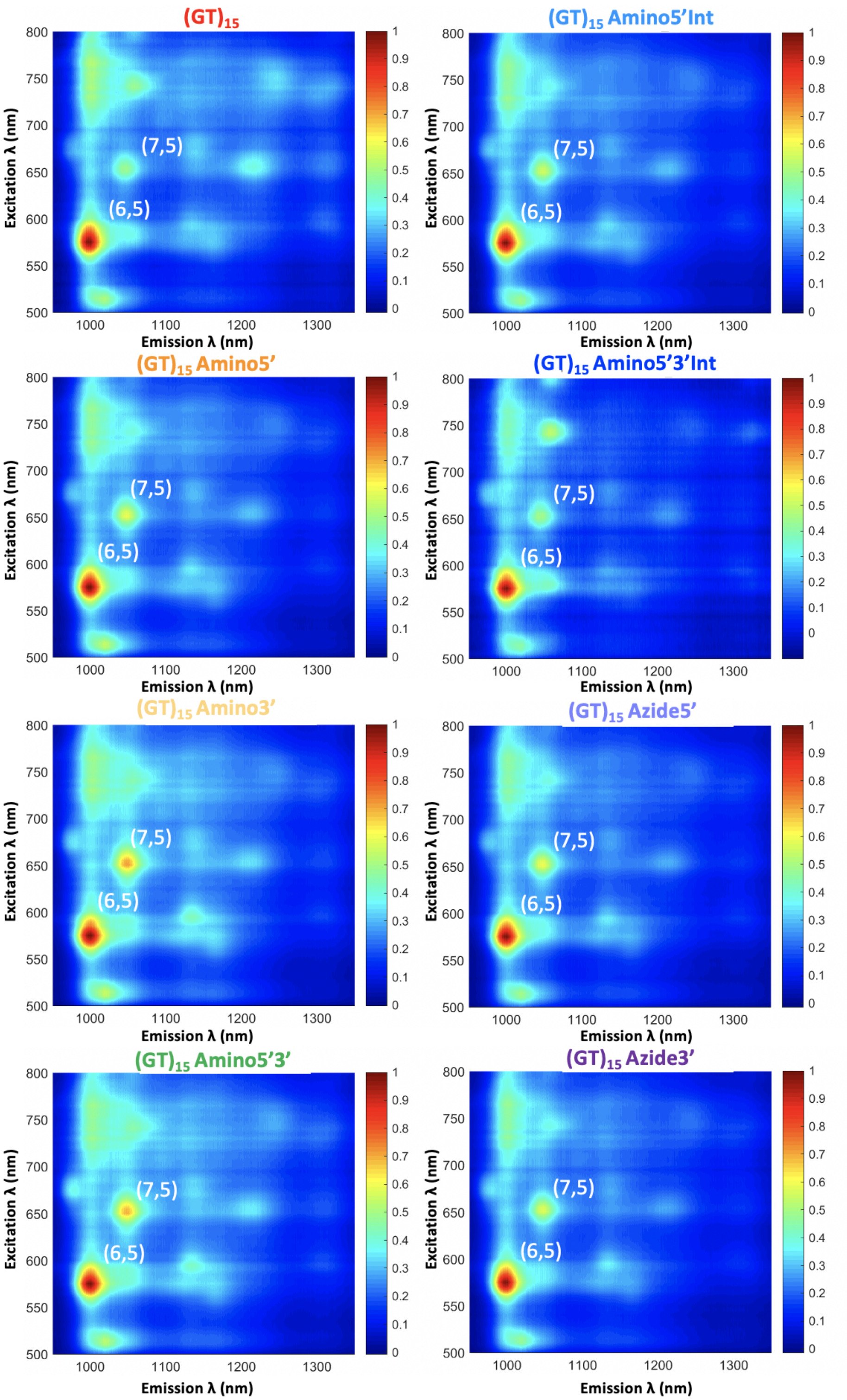


Figure 10: Photoluminescence excitation (PLE) maps of the modified and unmodified $(GT)_{15}$-SWCNT solutions prepared via direct sonication. The (6,5) and (7,5) chirality peaks are indicated in white. All fluorescence intensities were normalized to the maximum intensity in each plot.

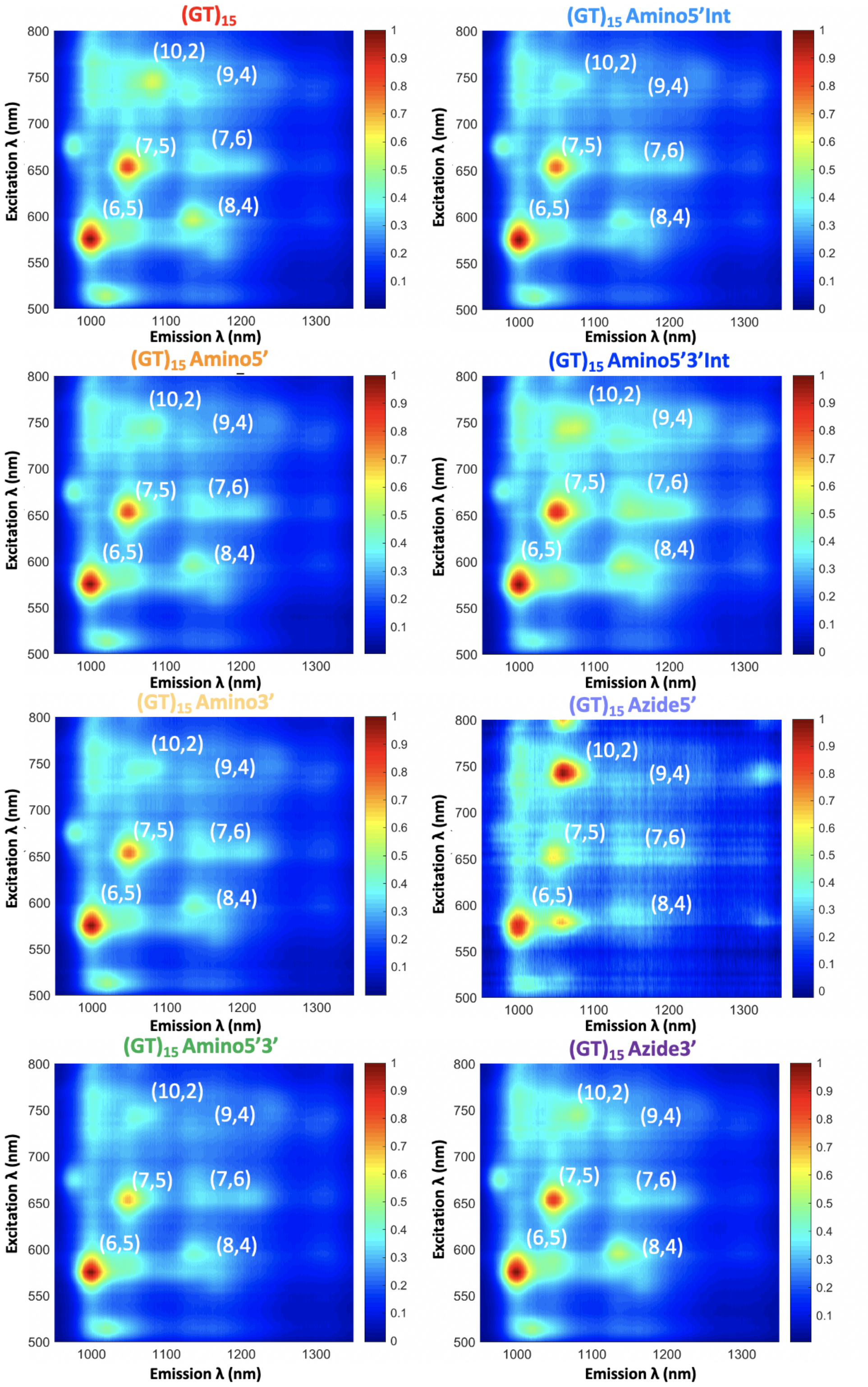


Figure 11: Photoluminescence excitation (PLE) maps of the modified and unmodified $(GT)_{15}$-SWCNT solutions prepared using MeOH assisted surfactant exchange. The (6,5) and (7,5) chirality peaks are indicated in white. All fluorescence intensities were normalized to the maximum intensity in each plot.

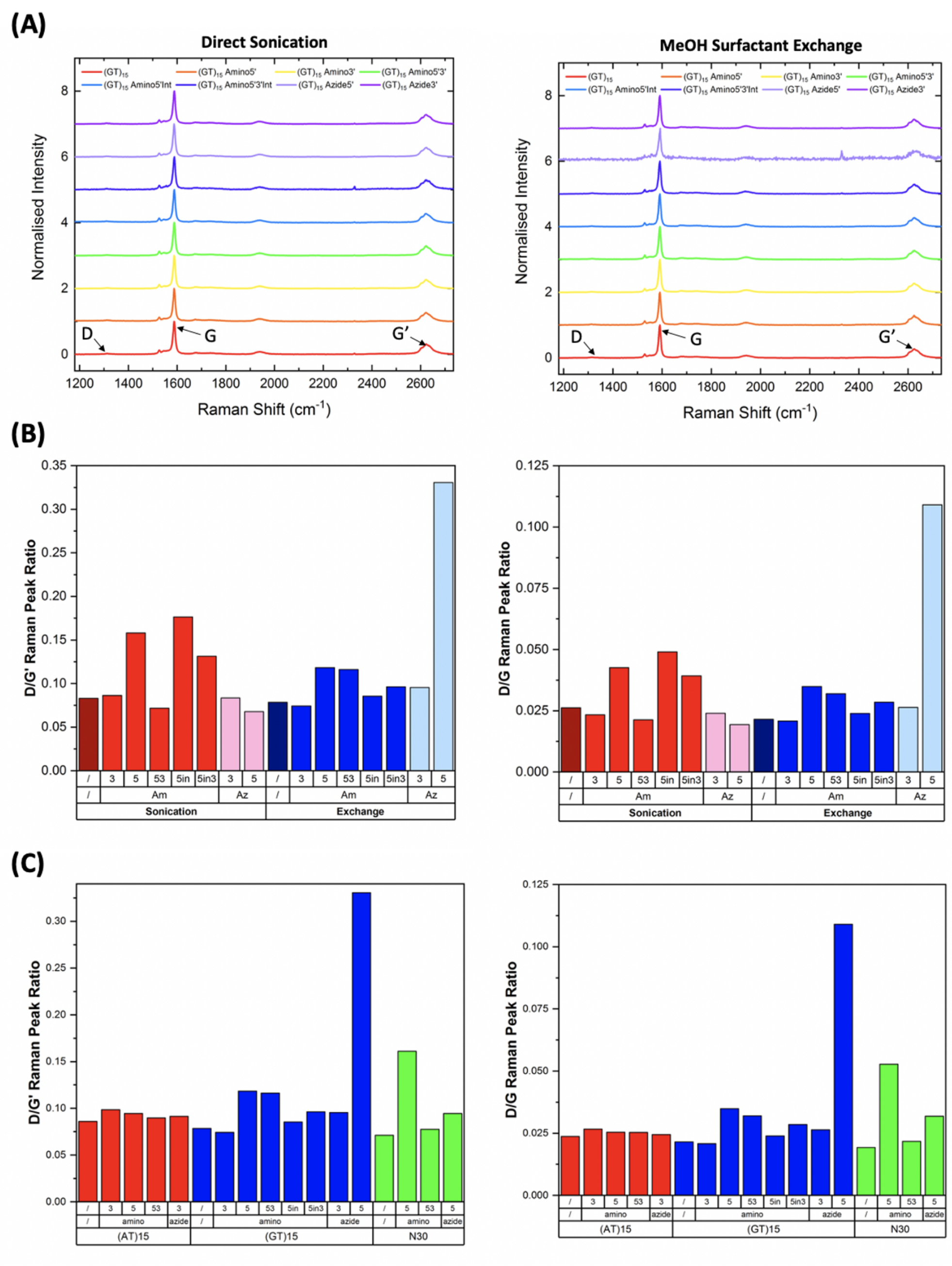


Figure 12: Raman characterisation of modified and unmodified DNA-SWCNTs prepared by direct sonication and MeOH assisted surfactant exchange. Nanotube surface defects provide the necessary symmetry breaking to give rise to the D-band in Raman spectra[1,2]. As a result, comparisons of the D/G' or D/G intensity ratios provide relative measures of defect density. The similarity of all Raman modes, specifically the D, G, and G' bands, for unmodified $(GT)_{15}$ samples prepared by both methods implies a common degree of disorder, suggesting that changes in the fluorescence intensity were not a result of increasing defect density on the sonicated samples. **(A)** Raman spectra for modified and unmodified $(GT)_{15}$-SWCNTs prepared by (**left**) direct sonication and (**right**) MeOH assisted surfactant exchange. Spectra were normalized to the G-band intensity and off-set for comparison. **(B)** Comparison of the (**left**) D/G' and (**right**) D/G peak ratio for the $(GT)_{15}$-SWCNT suspensions. **(C)** Comparison of the (**left**) D/G' and (**right**) D/G peak ratio for all DNA-SWCNT suspensions prepared using MeOH assisted surfactant exchange.

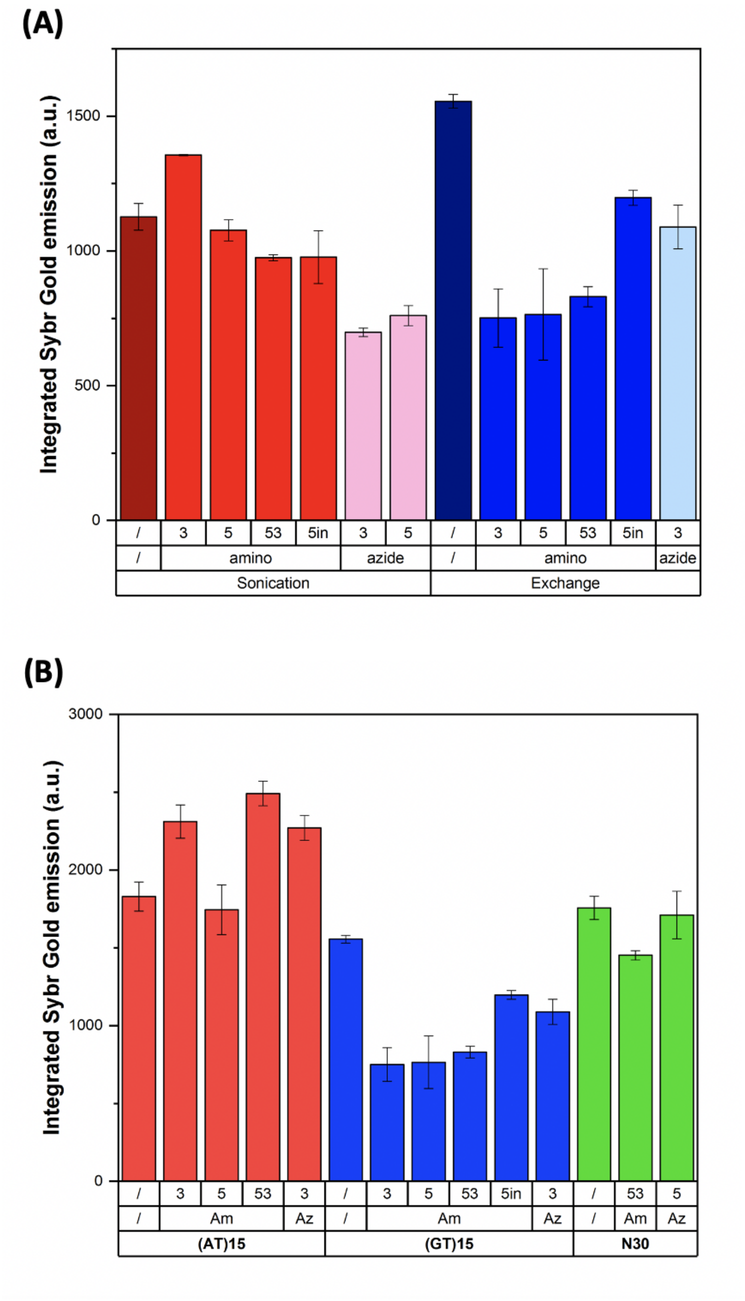


Figure 13: SYBR gold assay performed in order to calculate the amount of DNA adsorbed onto the surface of the nanotube for modified and unmodified **(A)** $(GT)_{15}$-SWCNTs prepared by direct sonication or MeOH assisted surfactant exchange and **(B)** DNA-SWCNTs prepared via MeOH assisted surfactant exchange. DNA was removed from the surface of the nanotube by phenol-chloroform isoamyl (PCI) extraction and subsequently precipitated by glycogen-assisted ethanol precipitation. Prior to DNA extraction, all DNA-SWCNT samples were rinsed using Amicon ultrafiltration devices to remove any unbound DNA that may be present in the solution. Higher integrated SYBR Gold emission values indicate higher concentrations of DNA. Error bars represent $1\sigma$ standard deviation ($n = 3$ technical replicates).

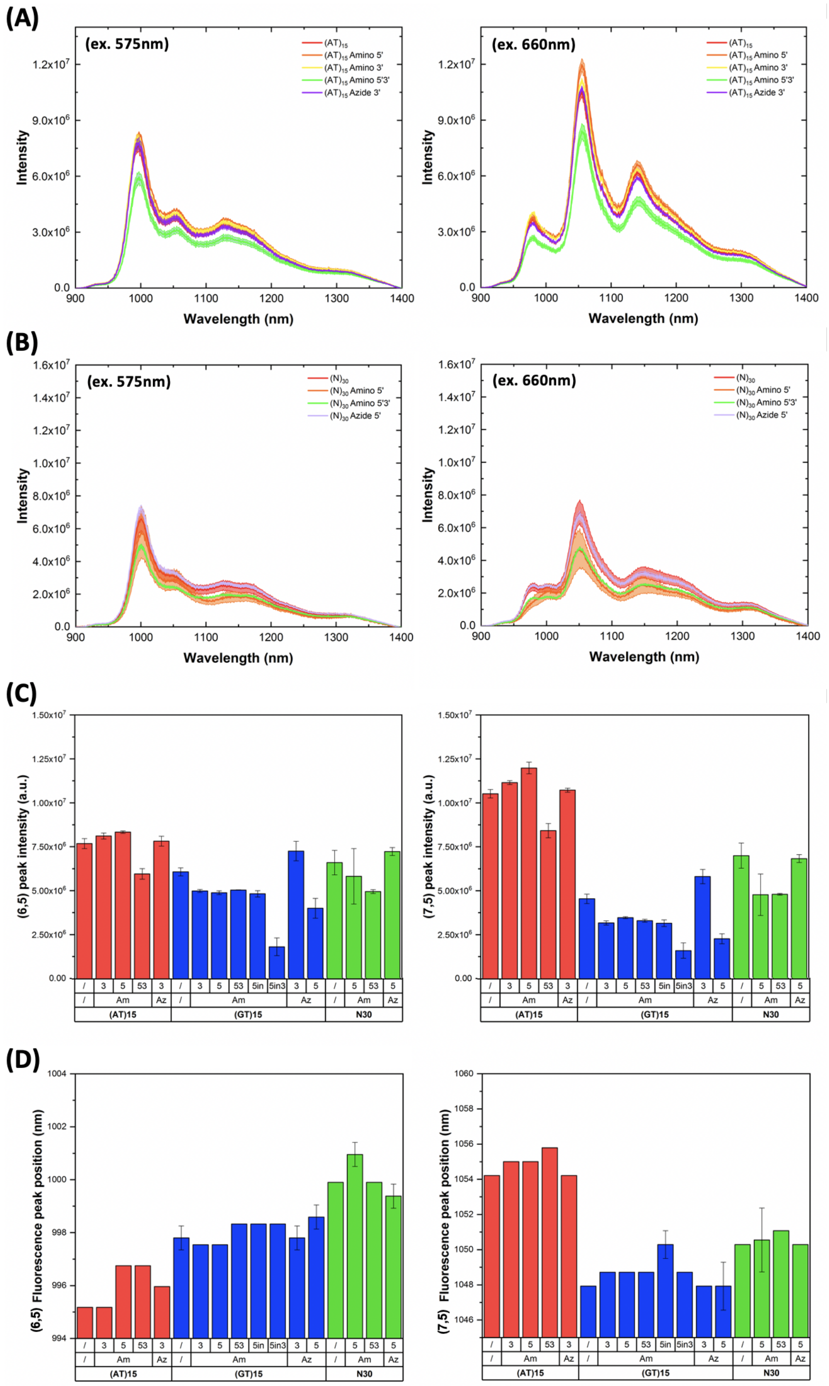


Figure 14: Fluorescence spectra of **(A)** $(AT)_{15}$- and **(B)** $(N)_{30}$-SWCNTs prepared using MeOH assisted surfactant exchange using both unmodified DNA (red line) and modified DNA sequences. For all spectra, the central line represents the average spectrum with the shaded regions representing $1\sigma$ standard deviation (n = 3 technical replicates) (excitation: 575 nm (**left**) and 660 nm (**right**). **(C)** Comparison of the absolute peak intensity of the (6,5) (excitation: 575 nm, (**left**)) and (7,5) chiralities (excitation: 660 nm, (**right**)) for modified and unmodified DNA-SWCNTs prepared using MeOH assisted surfactant exchange. Error bars represent $1\sigma$ standard deviation (n = 3 technical replicates). **(D)** Peak position of the (6,5) (excitation: 575 nm, (**left**)) and (7,5) chiralities (excitation: 660 nm, (**right**)) for modified and unmodified DNA-SWCNTs prepared using MeOH assisted surfactant exchange. Error bars represent $1\sigma$ standard deviation (n = 3 technical replicates).

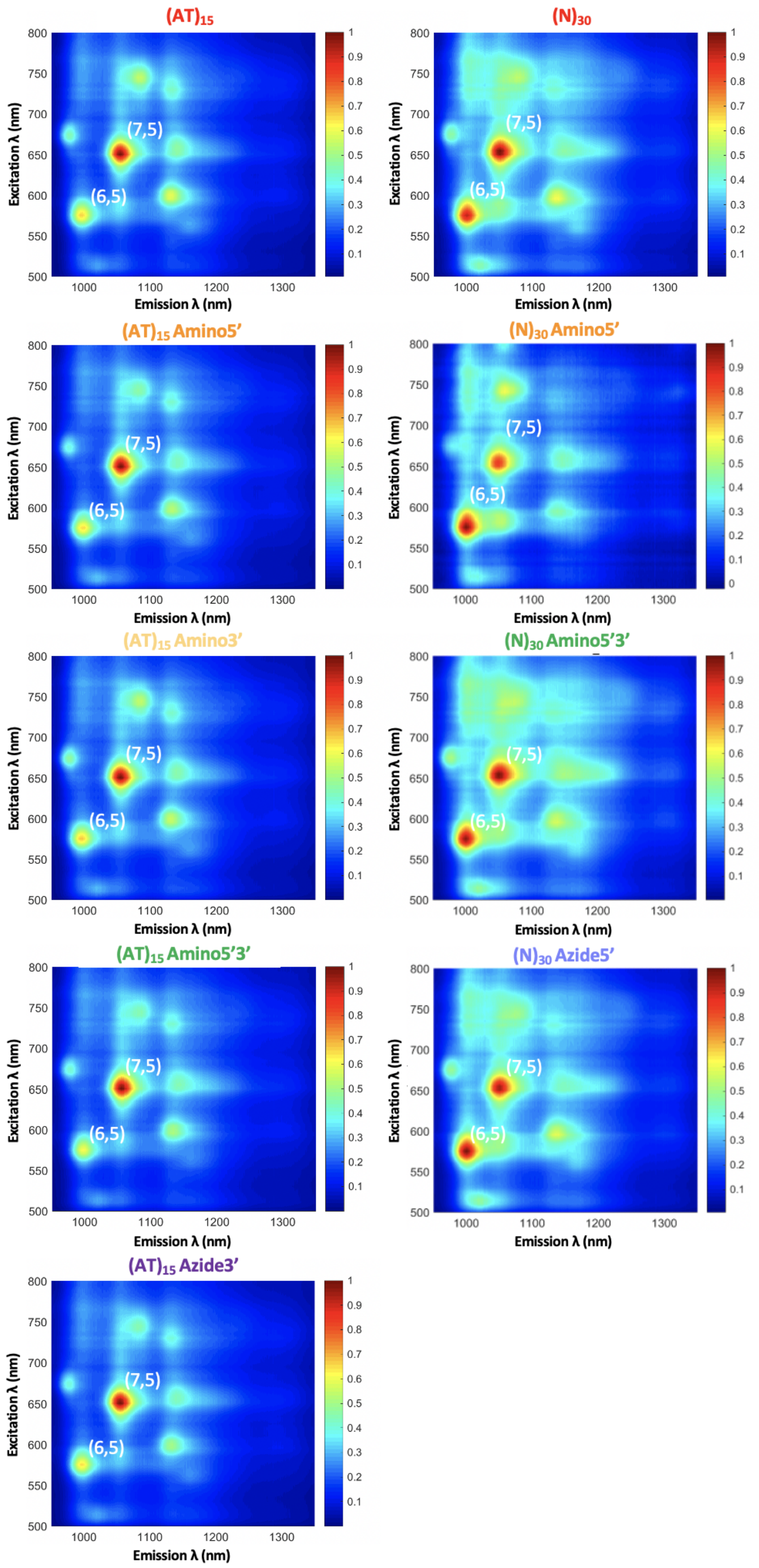


Figure 15: Photoluminescence excitation (PLE) maps of modified and unmodified $(AT)_{15}$- and $(N)_{30}$-SWCNTs prepared using MeOH assisted surfactant exchange. The (6,5) and (7,5) chirality peaks are indicated in white. All fluorescence intensities were normalized to the maximum intensity in each plot.

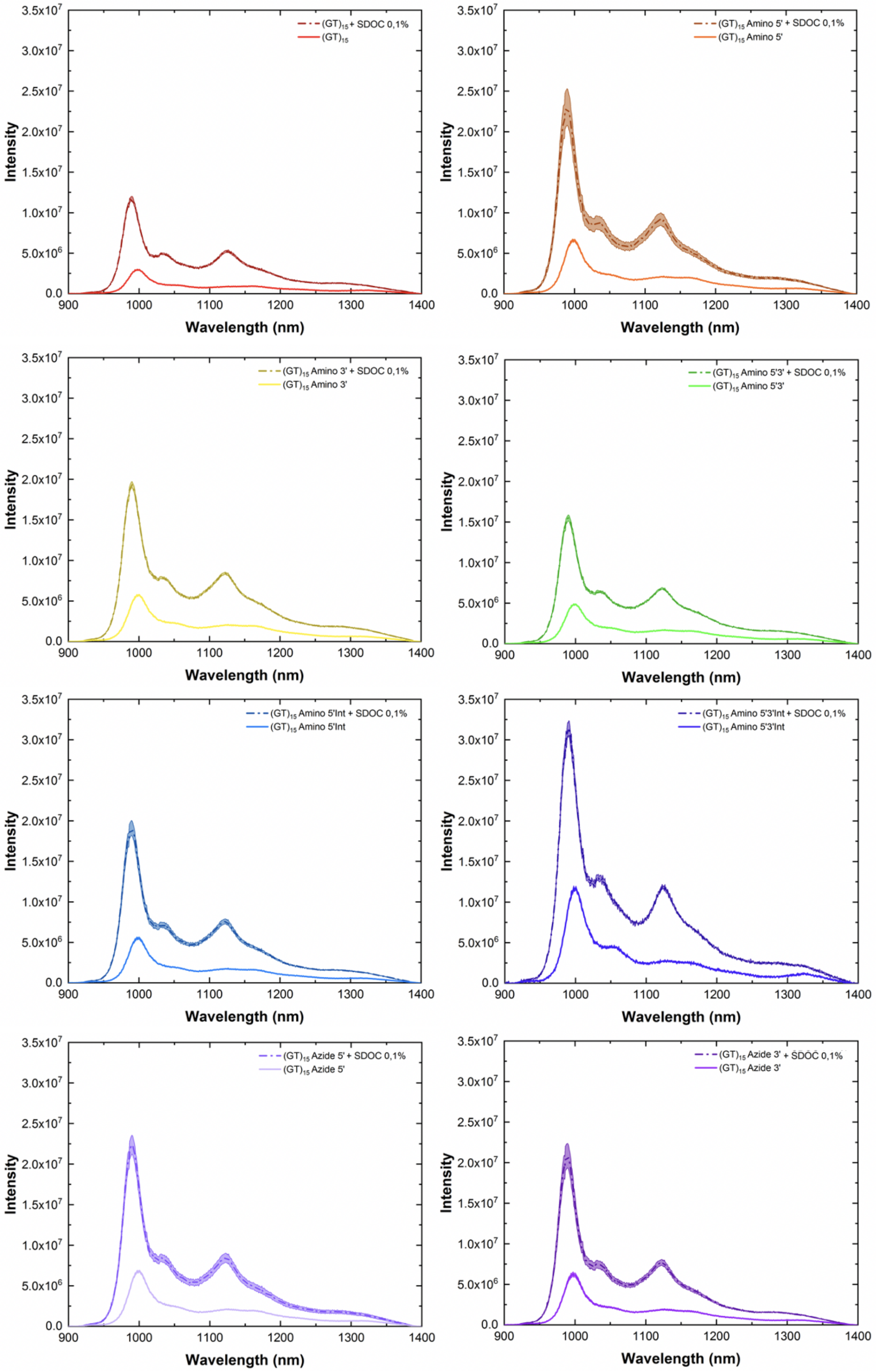


Figure 16: Fluorescence spectra of modified- and unmodified-$(GT)_{15}$-SWCNTs prepared via direct sonication before (solid light line) and after (dashed dark line) addition of SDOC (final concentration: 0.1%, excitation: 575 nm, incubation time: 10 min). For all spectra, the central line represents the average spectrum with the shaded regions representing $1\sigma$ standard deviation (n = 3 technical replicates).

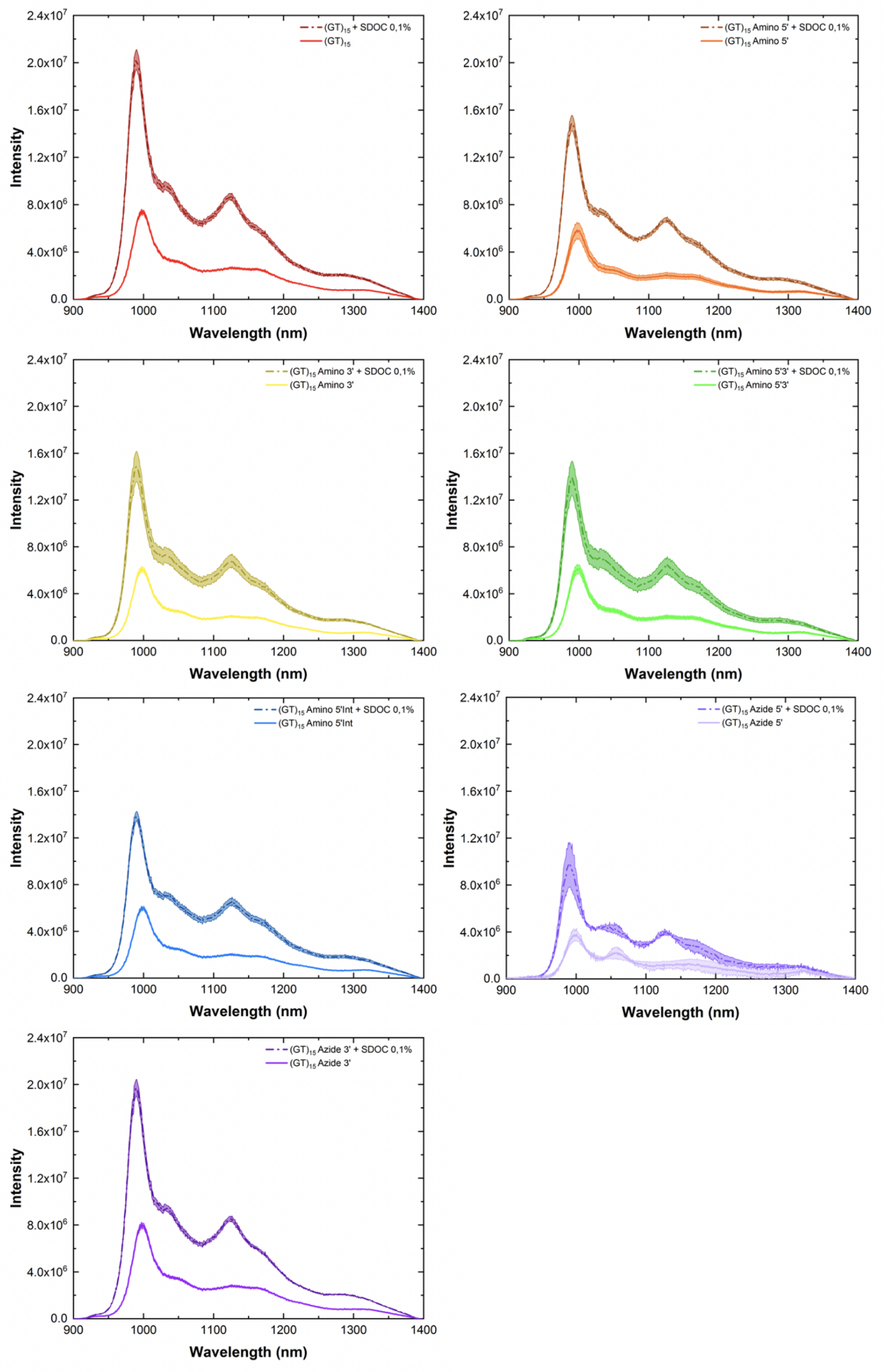


Figure 17: Fluorescence spectra of modified- and unmodified-(GT)$_{15}$-SWCNTs prepared using MeOH assisted surfactant exchange before (solid light line) and after (dashed dark line) addition of SDOC (final concentration: 0.1%, excitation: 575 nm, incubation time: 10 min). For all spectra, the central line represents the average spectrum with the shaded regions representing 1$\sigma$ standard deviation (n = 3 technical replicates).

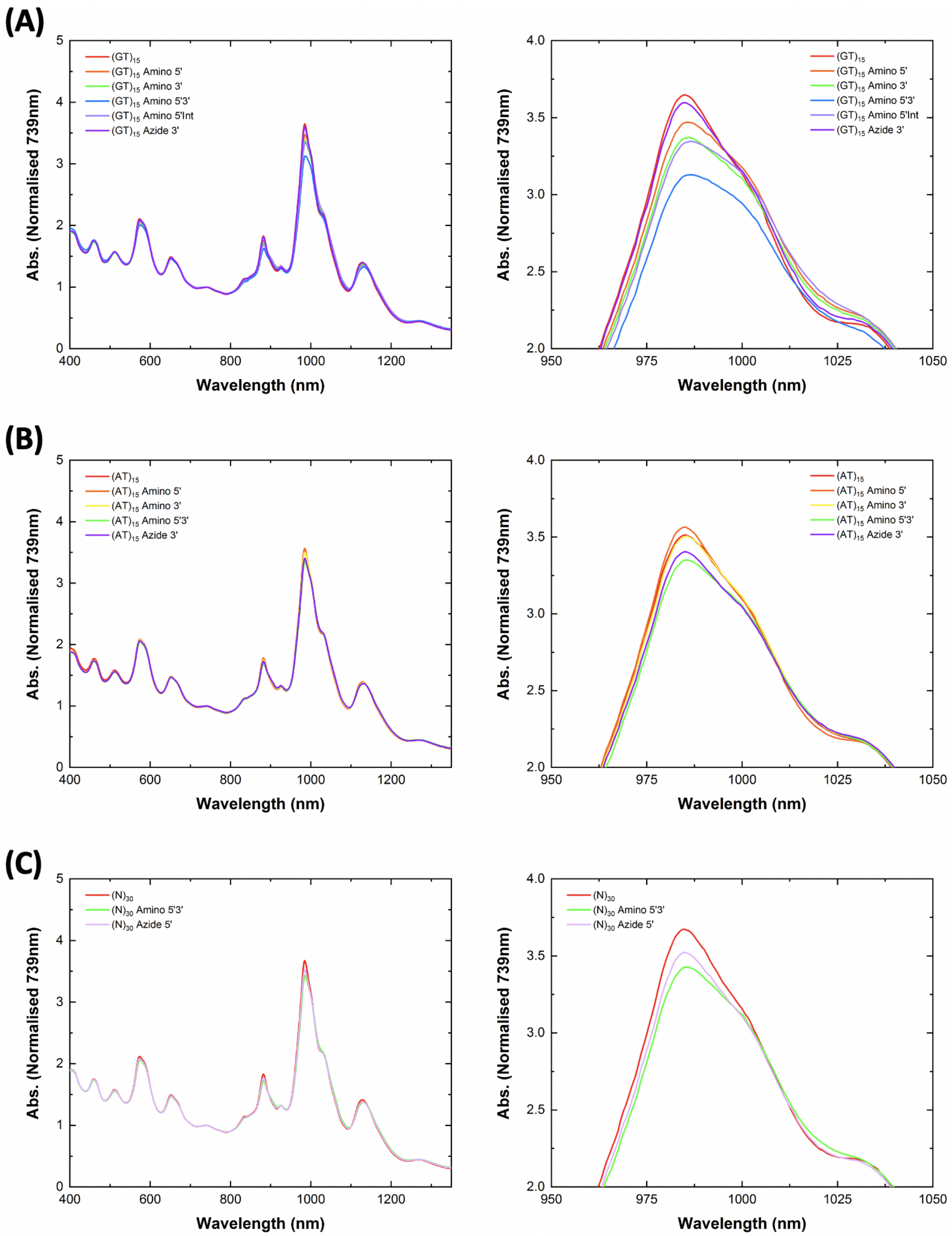


Figure 18: Absorbance spectra of modified and unmodified **(A)** $(GT)_{15}$-, **(B)** $(AT)_{15}$-, and **(C)** $(N)_{30}$-SWCNTs prepared using MeOH-assisted surfactant exchange following the addition of SDOC (final concentration: 0.1%). Spectra on the right are a zoom-in of the absorbance for the (6,5) chirality peak highlighting that even post SDOC replacement the absorbance did not overlap indicating that there were differences in the relative abundance of this chirality in the different suspensions. All spectra were normalized to the $Abs_{739nm}$ value to account for any differences in nanotube concentration.

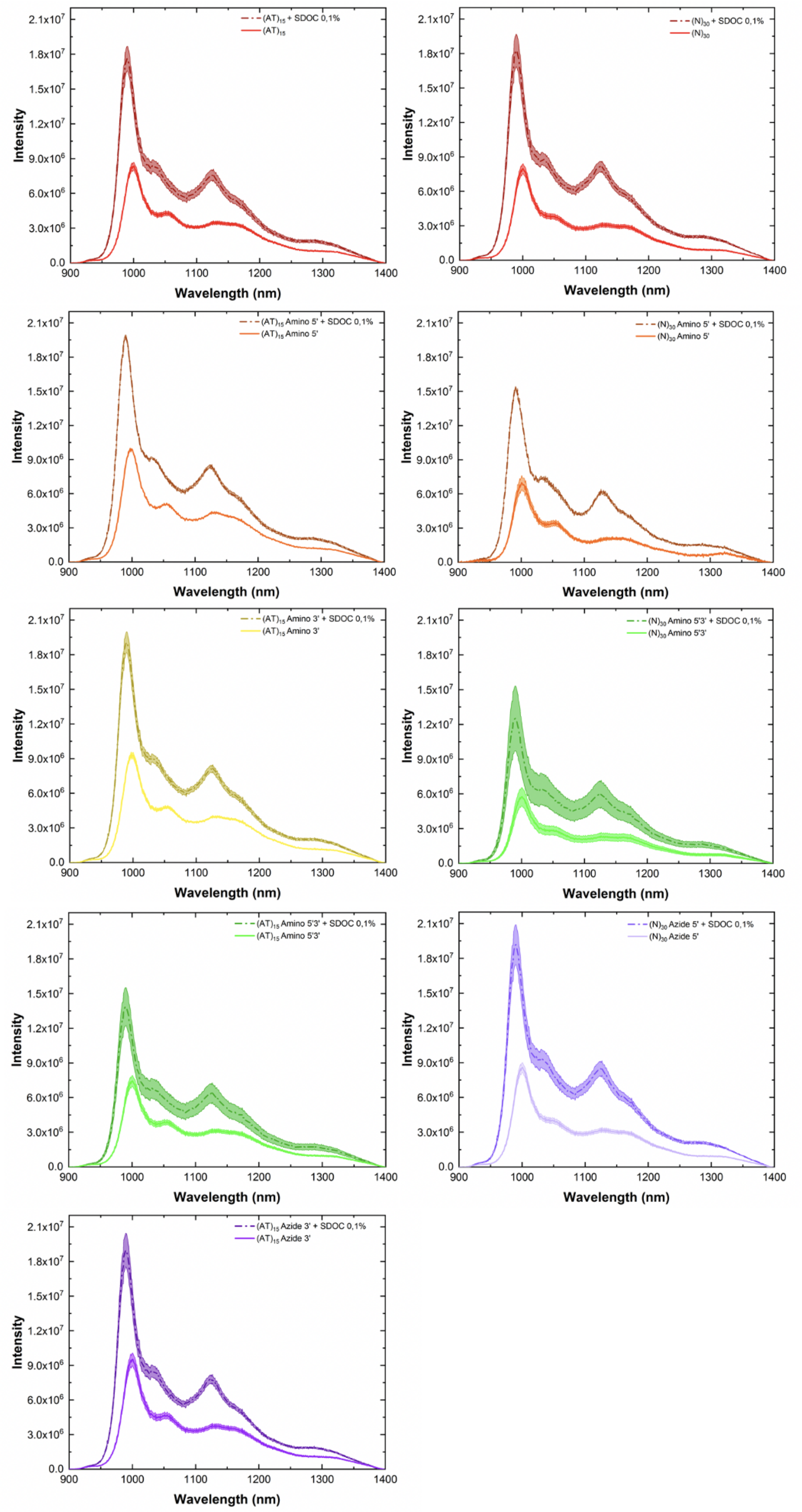


Figure 19: Fluorescence spectra of modified- and unmodified-$(AT)_{15}$-SWCNTs (**left**) and $(N)_{30}$-SWCNTs (**right**) prepared using MeOH-assisted surfactant exchange before (solid light line) and after (dashed dark line) addition of SDOC (final concentration: 0.1%, excitation: 575 nm, incubation period: 10 min). For all spectra, the central line represents the average spectrum with the shaded regions representing $1\sigma$ standard deviation (n = 3 technical replicates).

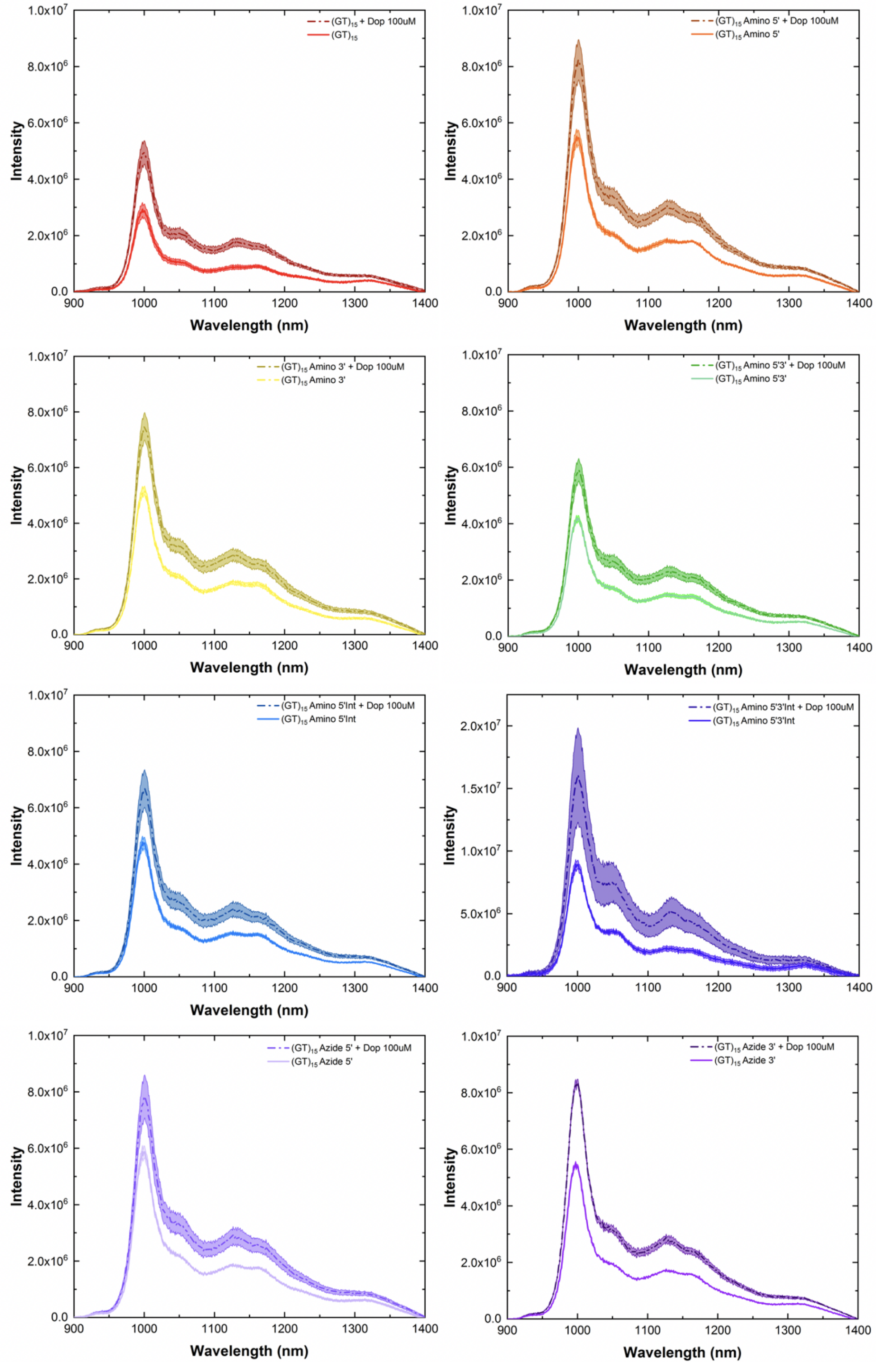


Figure 20: Fluorescence spectra of modified and unmodified $(GT)_{15}$-SWCNTs prepared via direct sonication before (solid light line) and after (dashed dark line) addition of dopamine (final concentration: 100 $\mu$M, excitation: 575 nm, incubation period: 10 min). For all spectra, the central line represents the average spectrum with the shaded regions representing $1\sigma$ standard deviation (n = 3 technical replicates).

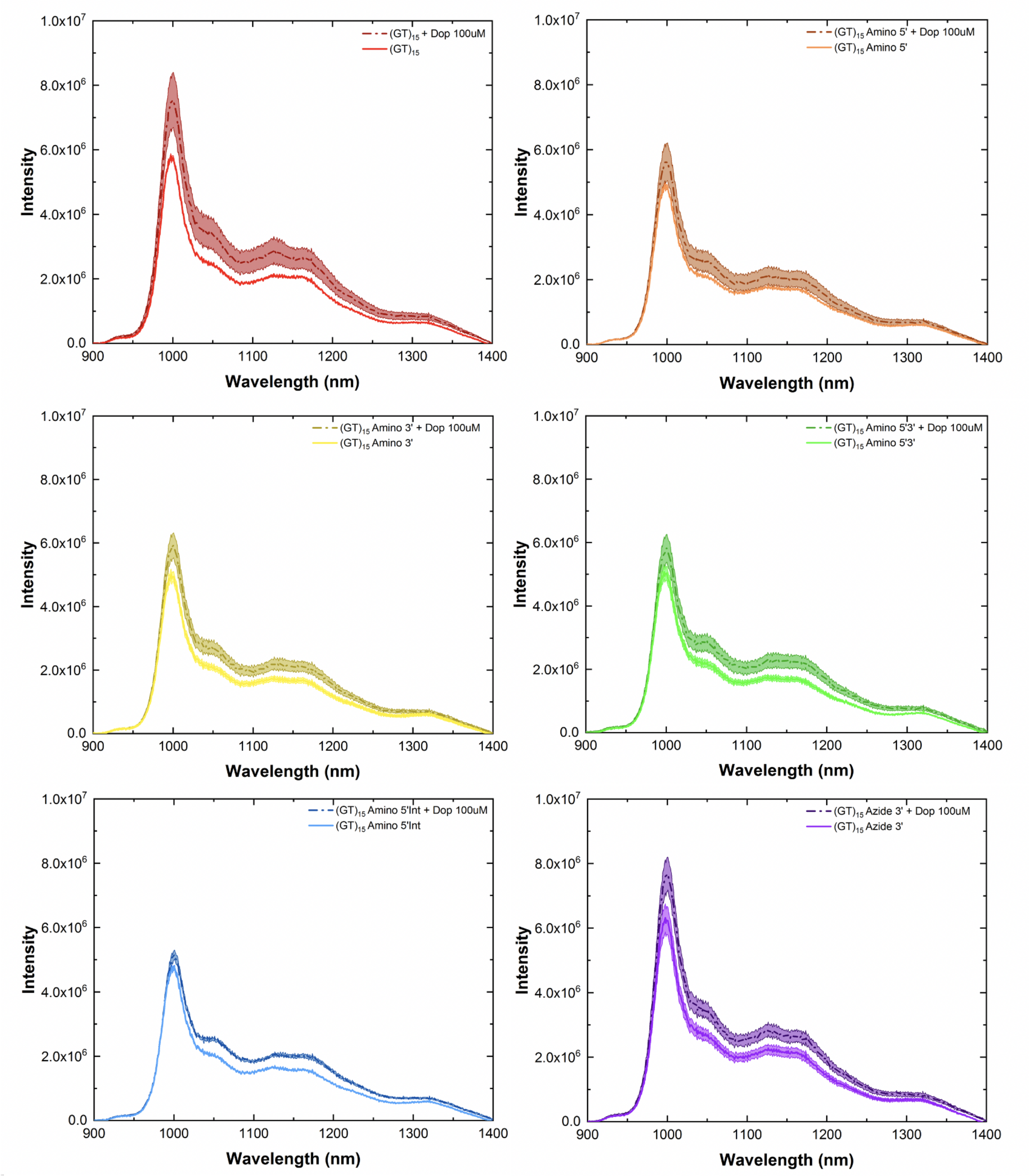


Figure 21: Fluorescence spectra of modified and unmodified $(GT)_{15}$-SWCNTs prepared using MeOH assisted surfactant exchange before (solid light line) and after (dashed dark line) addition of dopamine (final concentration: 100 $\mu$M, excitation: 575 nm, incubation period: 10 min). For all spectra, the central line represents the average spectrum with the shaded regions representing $1\sigma$ standard deviation (n = 3 technical replicates).

**Table 4:** Fluorescence intensity increase for modified and unmodified $(GT)_{15}$-SWCNTs following the addition of dopamine. Maximal fluorescence changes ($I_f$-$I_0/I_0$), % change, are presented alongside the absolute magnitude of the fluorescence increase ($I_f$-$I_0$) for comparison.

| **Direct Sonication** | | |
|---|---|---|
| **DNA Sequence** | % **Change** | **Absolute Increase** |
| $(GT)_{15}$-SWCNT | 69.1 ± 11.7 | 2.02 E+06 ± 0.48 E+06 |
| Amino3'-$(GT)_{15}$-SWCNT | 44.4 ± 7.2 | 2.30 E+06 ± 0.52 E+06 |
| Amino5'-$(GT)_{15}$-SWCNT | 49.4 ± 9.9 | 2.73 E+06 ± 0.76 E+06 |
| Amino5'3'-$(GT)_{15}$-SWCNT | 40.8 ± 7.1 | 1.71 E+06 ± 0.41 E+06 |
| Amino5'Int-$(GT)_{15}$-SWCNT | 39.7 ± 10.9 | 1.90 E+06 ± 0.70 E+06 |
| Azide3'-$(GT)_{15}$-SWCNT | 51.6 ± 1.7 | 2.85 E+06 ± 0.12 E+06 |

| **MeOH-assisted Surfactant Exchange** | | |
|---|---|---|
| **DNA Sequence** | % **Change** | **Magnitude of Increase** |
| $(GT)_{15}$-SWCNT | 30.1 ± 11.4 | 1.75 E+06 ± 0.86 E+06 |
| Amino3'-$(GT)_{15}$-SWCNT | 18.3 ± 7.4 | 9.18 E+05 ± 4.27 E+05 |
| Amino5'-$(GT)_{15}$-SWCNT | 14.5 ± 10.3 | 7.14 E+05 ± 5.82 E+05 |
| Amino5'3'-$(GT)_{15}$-SWCNT | 14.9 ± 8.5 | 7.57 E+05 ± 4.80 E+05 |
| Amino5'Int-$(GT)_{15}$-SWCNT | 8.4 ± 3.1 | 4.02 E+05 ± 1.57 E+05 |
| Azide3'-$(GT)_{15}$-SWCNT | 21.7 ± 9.5 | 1.37 E+06 ± 0.67 E+06 |

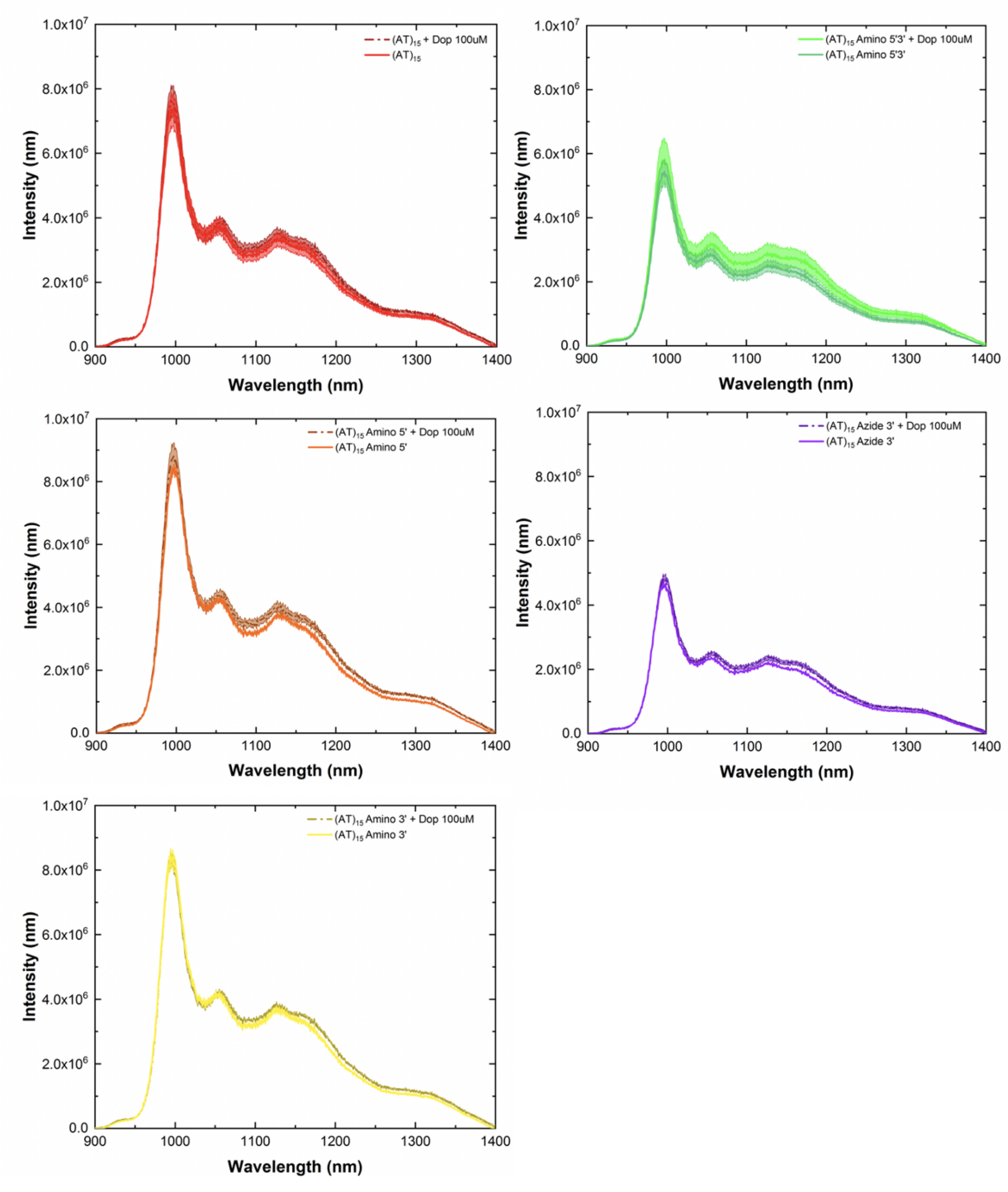


Figure 22: Fluorescence spectra of modified and unmodified $(AT)_{15}$-SWCNTs prepared using MeOH assisted surfactant exchange before (solid light line) and after (dashed dark line) addition of dopamine (final concentration: 100 $\mu$M, excitation: 575 nm, incubation period: 10 min). For all spectra, the central line represents the average spectrum with the shaded regions representing $1\sigma$ standard deviation (n = 3 technical replicates).

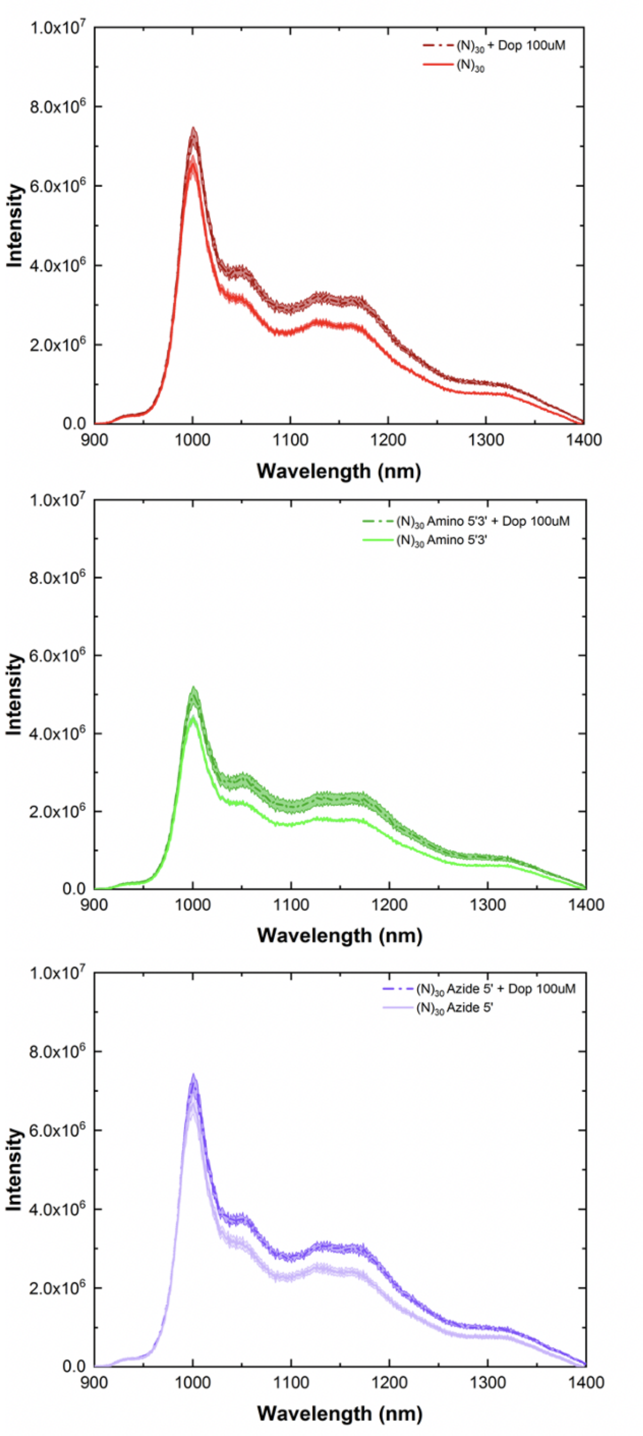


Figure 23: Fluorescence spectra of modified and unmodified (N)$_{30}$-SWCNTs prepared using MeOH assisted surfactant exchange before (solid light line) and after (dashed dark line) addition of dopamine (final concentration: 100 $\mu$M, excitation: 575 nm, incubation period: 10 min). For all spectra, the central line represents the average spectrum with the shaded regions representing 1$\sigma$ standard deviation (n = 3 technical replicates)..

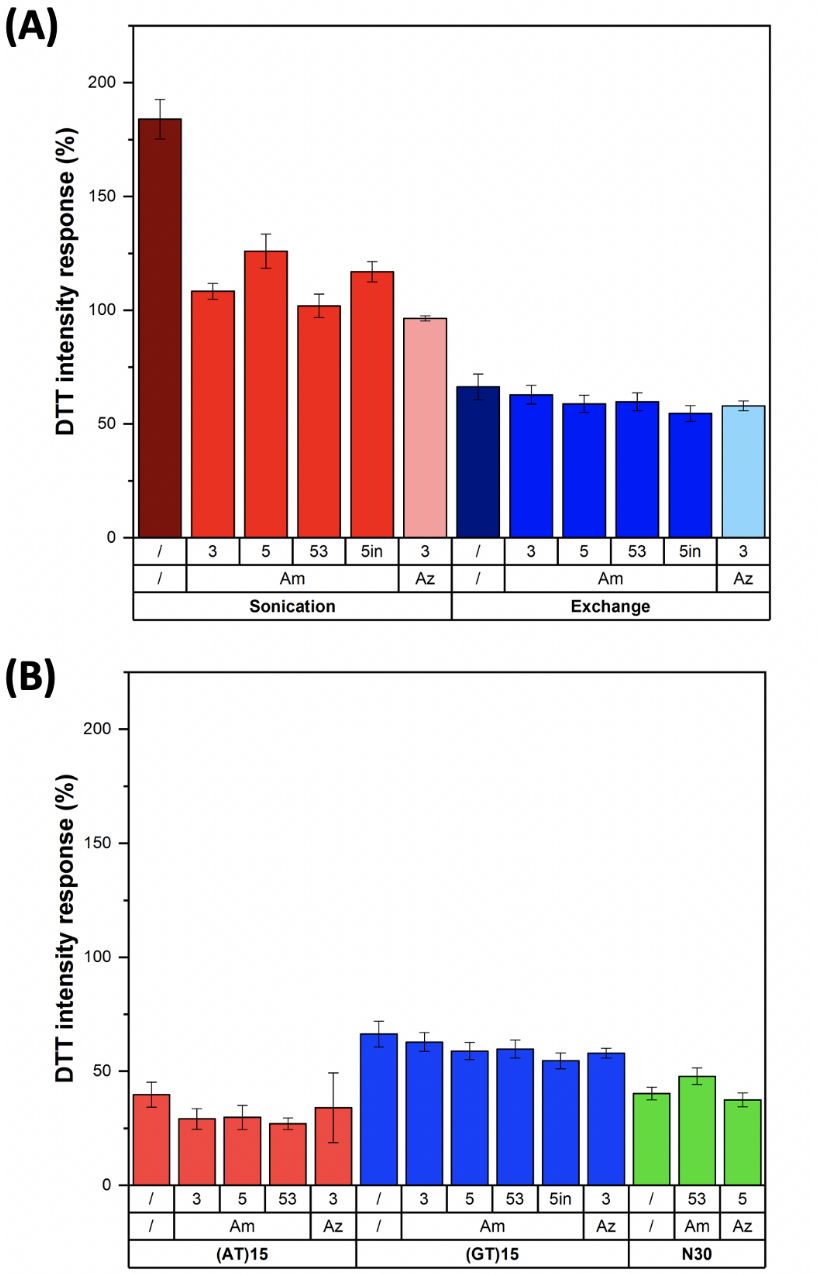


Figure 24: Intensity response following the addition of DTT. Comparison of the response for modified and unmodified **(A)** $(GT)_{15}$-SWCNTs prepared by direct sonication and MeOH-assisted surfactant exchange and **(B)** DNA-SWCNTs prepared using MeOH assisted surfactant exchange towards DTT. Intensity changes were calculated as $I_f$-$I_0/I_0$ for the (6,5) chirality peak (final concentration: 10 mM, excitation: 575 nm, incubation period: 10 min). Error bars represent $1\sigma$ standard deviation (n = 3 technical replicates).

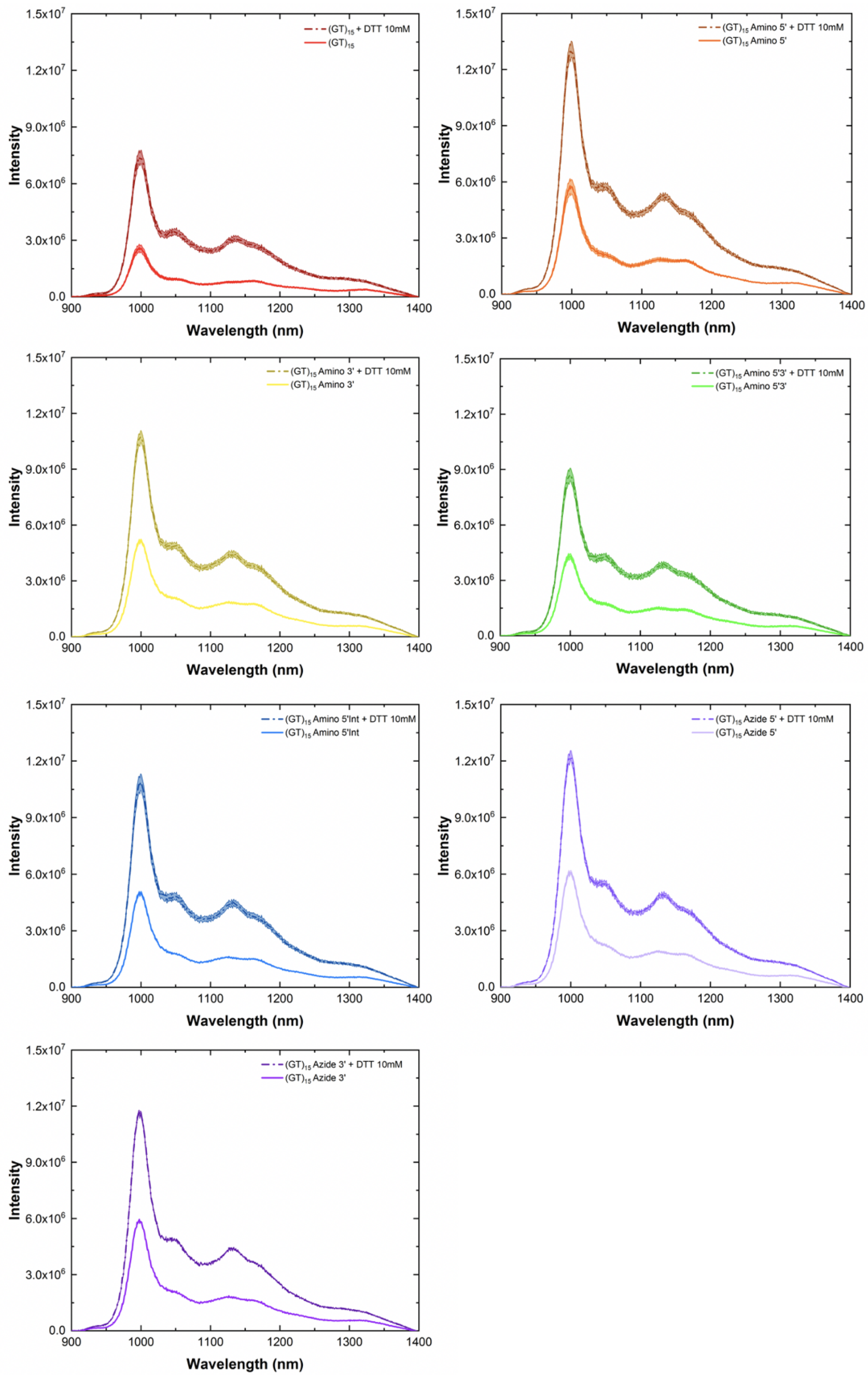


Figure 25: Fluorescence spectra of modified and unmodified (GT)$_{15}$-SWCNTs prepared by direct sonication before (solid light line) and after (dashed dark line) addition of DTT (final concentration: 10 mM, excitation: 575 nm, incubation period: 10 min). For all spectra, the central line represents the average spectrum with the shaded regions representing $1\sigma$ standard deviation (n = 3 technical replicates).

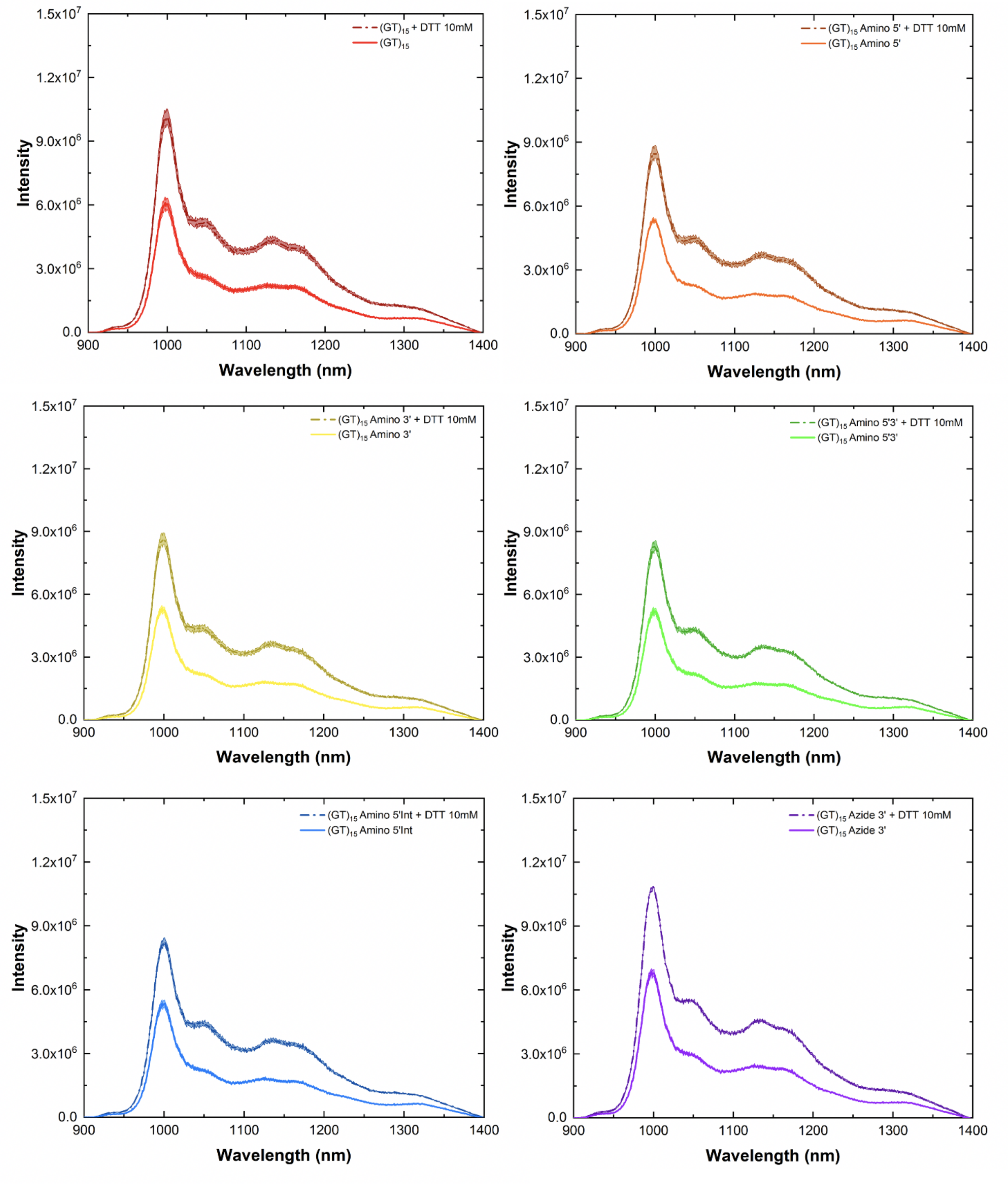


Figure 26: Fluorescence spectra of modified and unmodified $(GT)_{15}$-SWCNTs prepared using MeOH assisted surfactant exchange before (solid light line) and after (dashed dark line) addition of DTT (final concentration: 10 mM, excitation: 575 nm, incubation period: 10 min). For all spectra, the central line represents the average spectrum with the shaded regions representing $1\sigma$ standard deviation (n = 3 technical replicates).

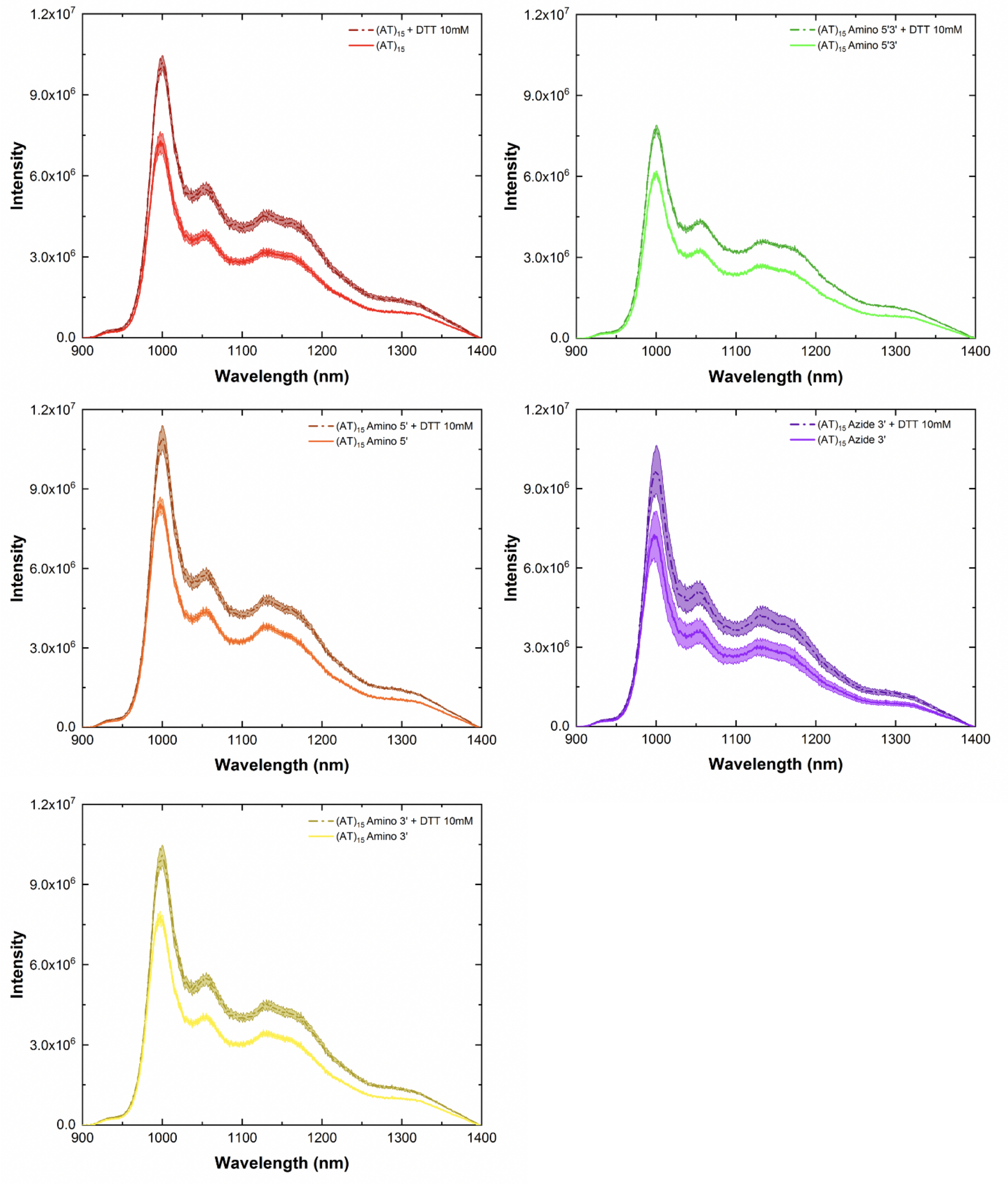


Figure 27: Fluorescence spectra of modified and unmodified $(AT)_{15}$-SWCNTs prepared using MeOH assisted surfactant exchange before (solid light line) and after (dashed dark line) addition of DTT (final concentration: 10 mM, excitation: 575 nm, incubation period: 10 min). For all spectra, the central line represents the average spectrum with the shaded regions representing $1\sigma$ standard deviation (n = 3 technical replicates).

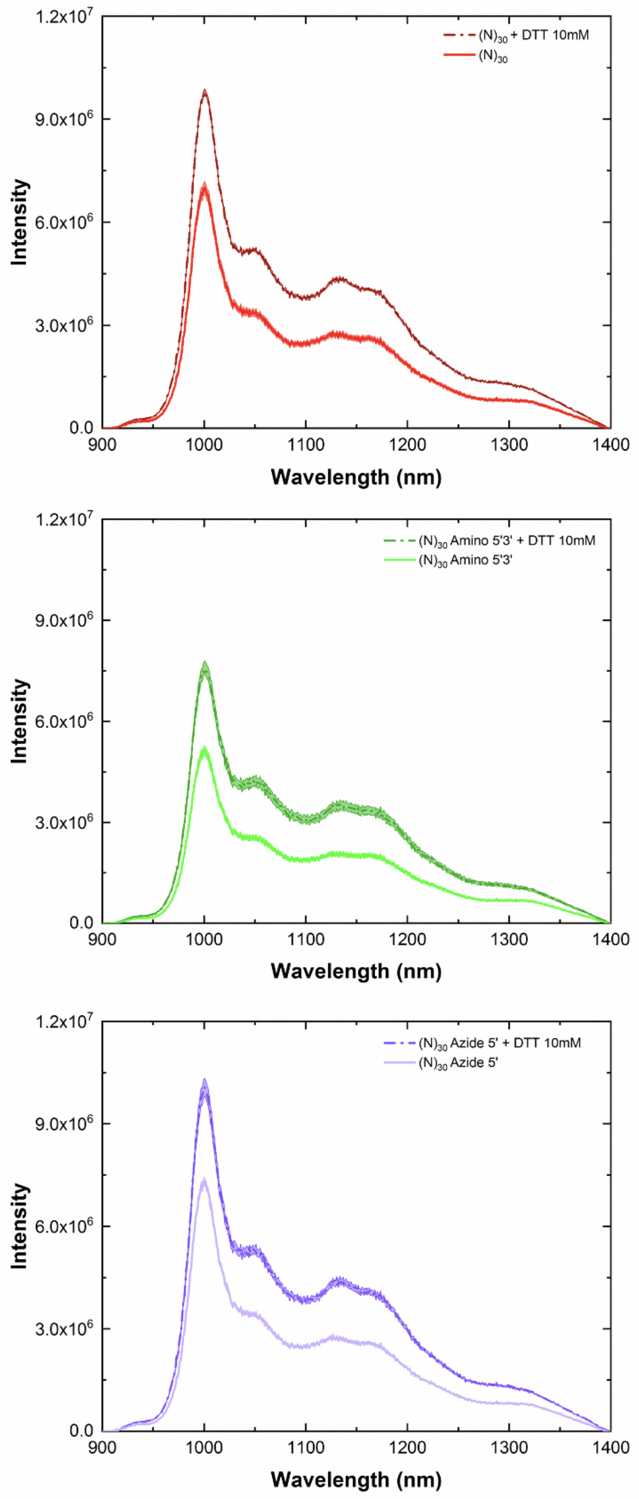


Figure 28: Fluorescence spectra of modified and unmodified $(N)_{30}$-SWCNTs prepared using MeOH assisted surfactant exchange before (solid light line) and after (dashed dark line) addition of DTT (final concentration: 10 mM, excitation: 575 nm, incubation period: 10 min). For all spectra, the central line represents the average spectrum with the shaded regions representing $1\sigma$ standard deviation (n = 3 technical replicates).

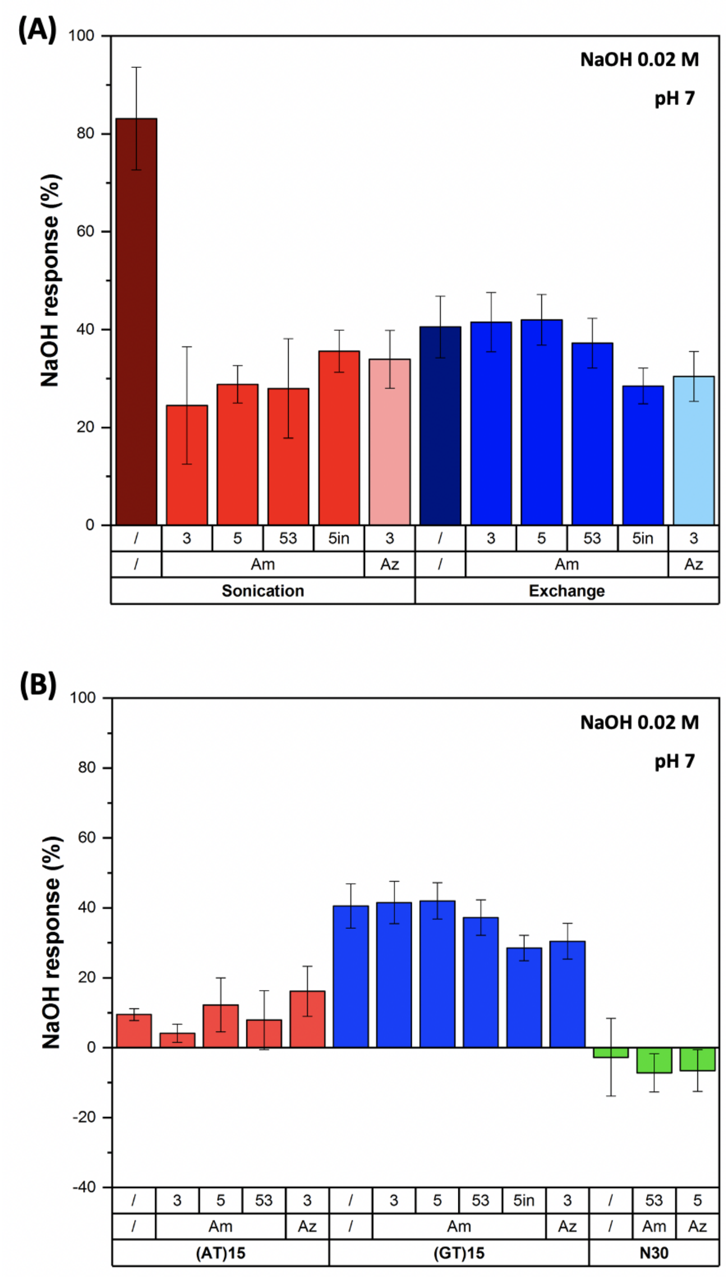


Figure 29: Intensity response following the addition of NaOH 0.02 M to increase the pH of the solution to pH 7. Comparison of the response for modified and unmodified **(A)** $(GT)_{15}$-SWCNTs prepared by direct sonication and MeOH assisted surfactant exchange and **(B)** DNA-SWCNTs prepared using MeOH-assisted surfactant exchange. Intensity changes were calculated as $I_f$-$I_0/I_0$ for the (6,5) chirality peak (final pH: 7, excitation: 575 nm, incubation period: 10 min). Error bars represent $1\sigma$ standard deviation (n = 3 technical replicates).

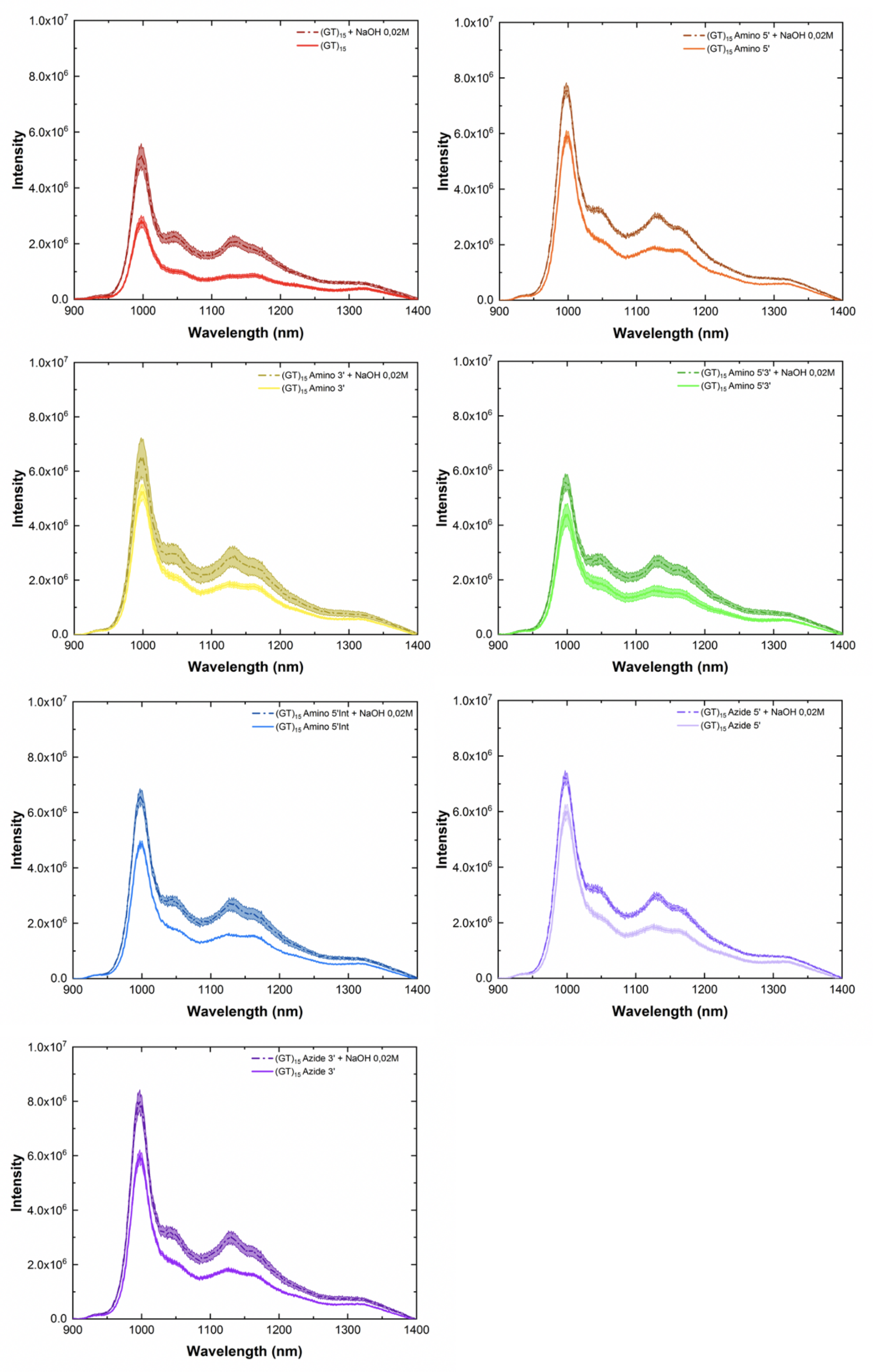


Figure 30: Fluorescence spectra of modified and unmodified $(GT)_{15}$-SWCNTs prepared by direct sonication before (solid light line) and after (dashed dark line) addition of NaOH 0.02 M (final pH: 7, excitation: 575 nm, incubation period: 10 min). For all spectra, the central line represents the average spectrum with the shaded regions representing $1\sigma$ standard deviation (n = 3 technical replicates).

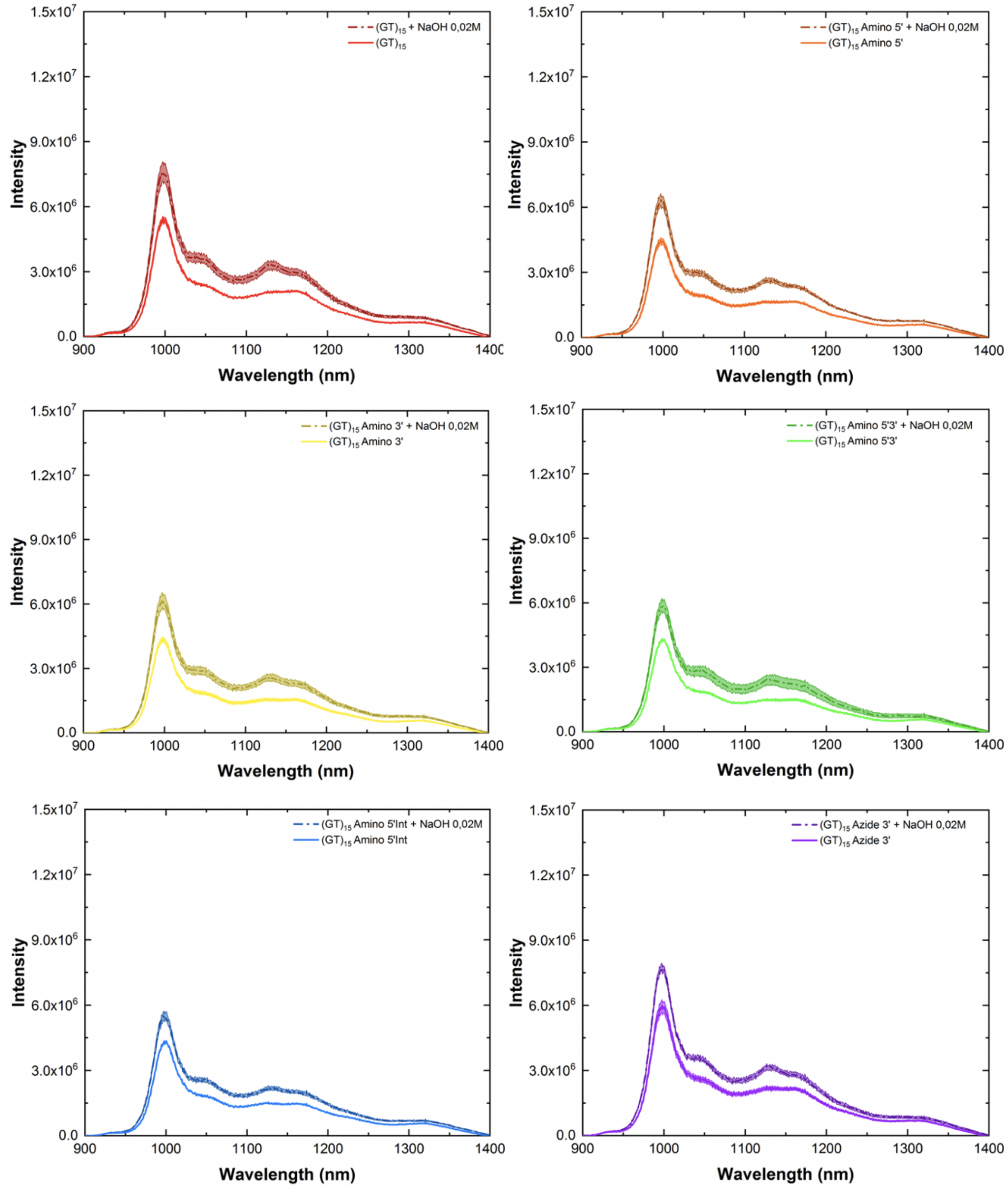


Figure 31: Fluorescence spectra of modified and unmodified $(GT)_{15}$-SWCNTs prepared using MeOH assisted surfactant exchange before (solid light line) and after (dashed dark line) addition of NaOH 0.02 M (final pH: 7, excitation: 575 nm, incubation period: 10 min). For all spectra, the central line represents the average spectrum with the shaded regions representing $1\sigma$ standard deviation (n = 3 technical replicates).

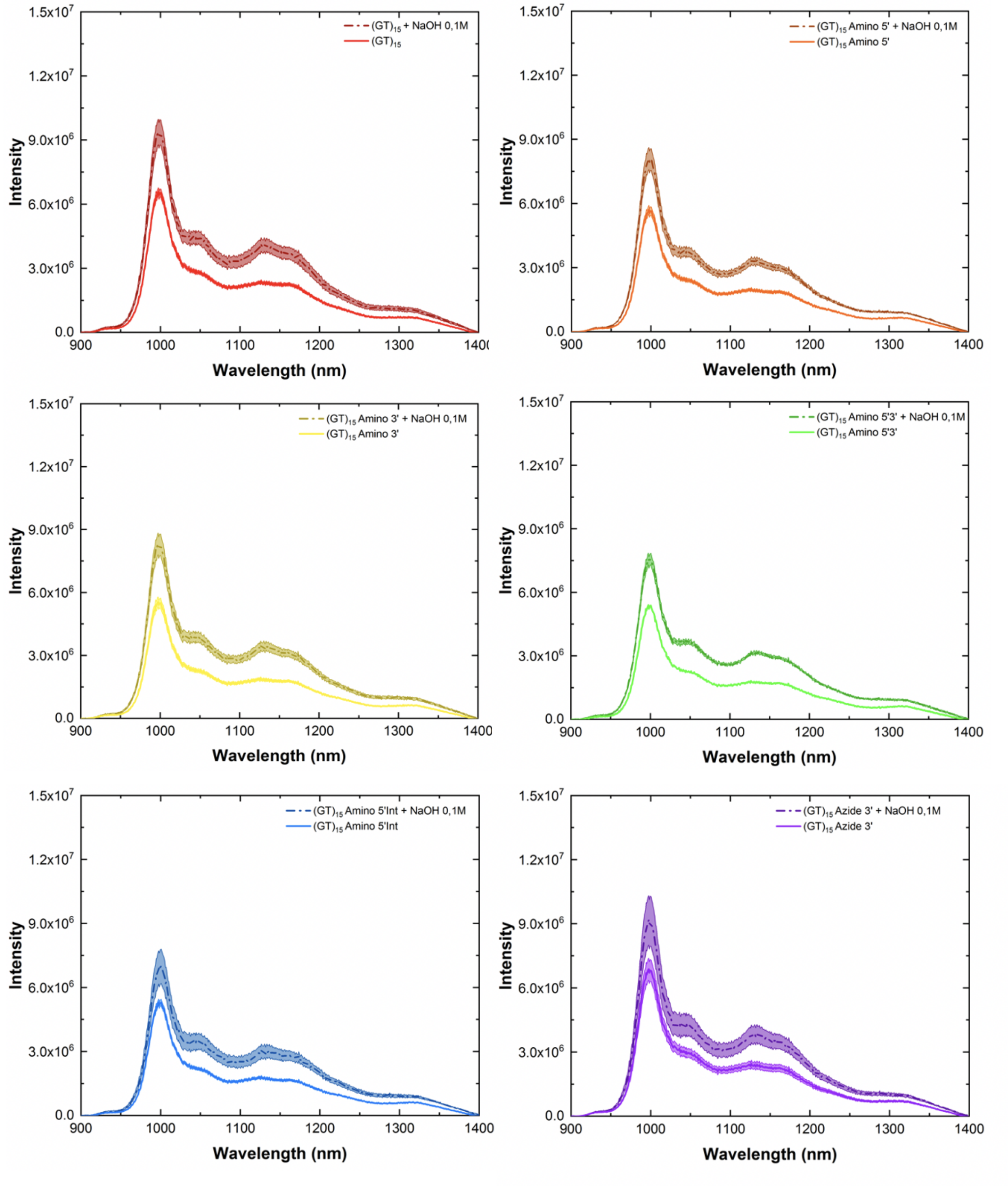


Figure 32: Fluorescence spectra of modified and unmodified $(GT)_{15}$-SWCNTs prepared using MeOH assisted surfactant exchange before (solid light line) and after (dashed dark line) addition of NaOH 0.1 M (final pH: 9, excitation: 575 nm, incubation period: 10 min). For all spectra, the central line represents the average spectrum with the shaded regions representing $1\sigma$ standard deviation (n = 3 technical replicates).

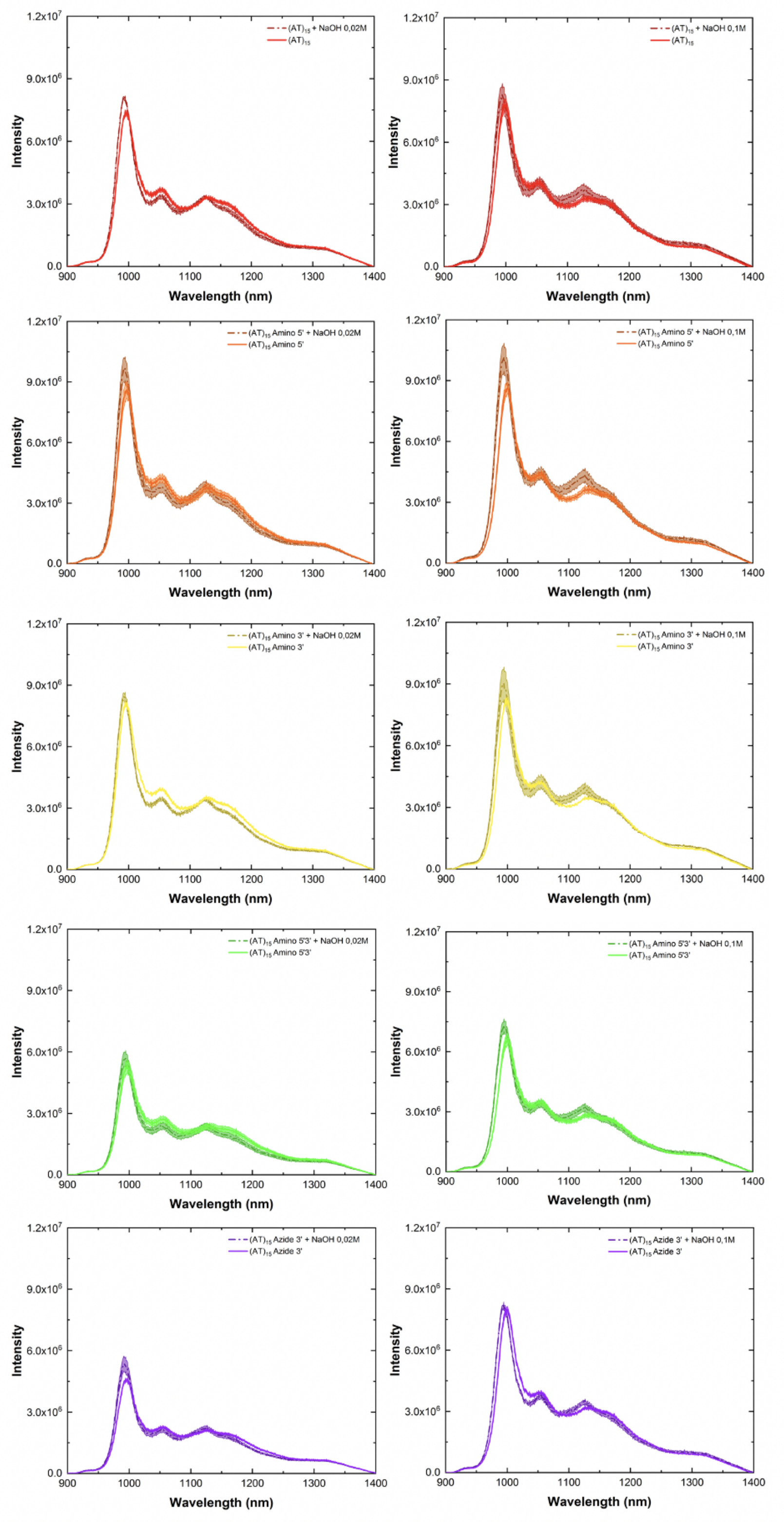


Figure 33: Fluorescence spectra of modified and unmodified $(AT)_{15}$-SWCNTs prepared using MeOH assisted surfactant exchange before (solid light line) and after (dashed dark line) addition of (**left**) NaOH 0.02 M (final pH: 7) and (**right**) NaOH 0.1 M (final pH: 9) (excitation: 575 nm, incubation period: 10 min). For all spectra, the central line represents the average spectrum with the shaded regions representing $1\sigma$ standard deviation (n = 3 technical replicates).

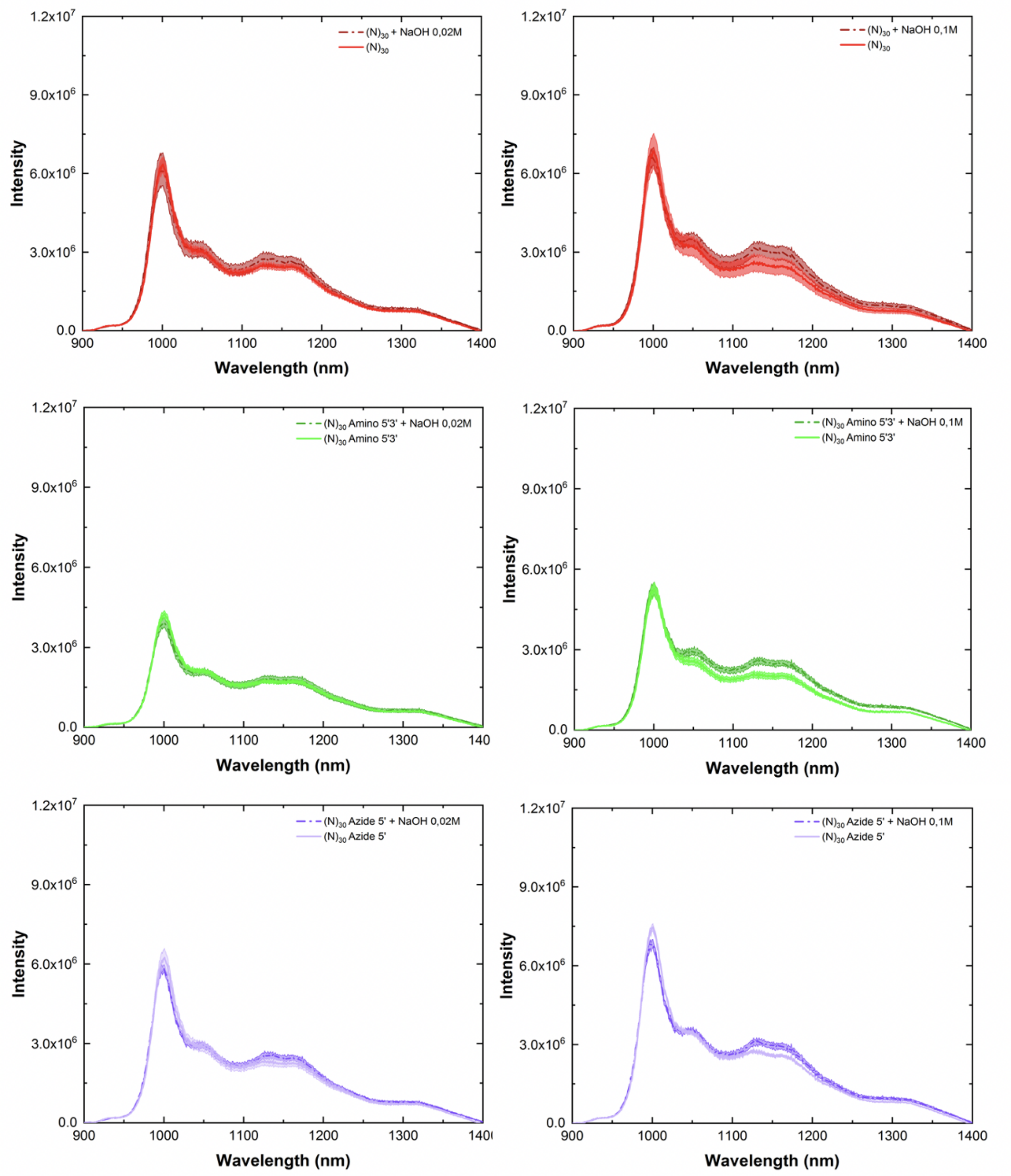


Figure 34: Fluorescence spectra of modified and unmodified $(N)_{30}$-SWCNTs prepared using MeOH assisted surfactant exchange before (solid light line) and after (dashed dark line) addition of (**left**) NaOH 0.02 M (final pH: 7) and (**right**) NaOH 0.1 M (final pH: 9) (excitation: 575 nm, incubation period: 10 min). For all spectra, the central line represents the average spectrum with the shaded regions representing $1\sigma$ standard deviation (n = 3 technical replicates).

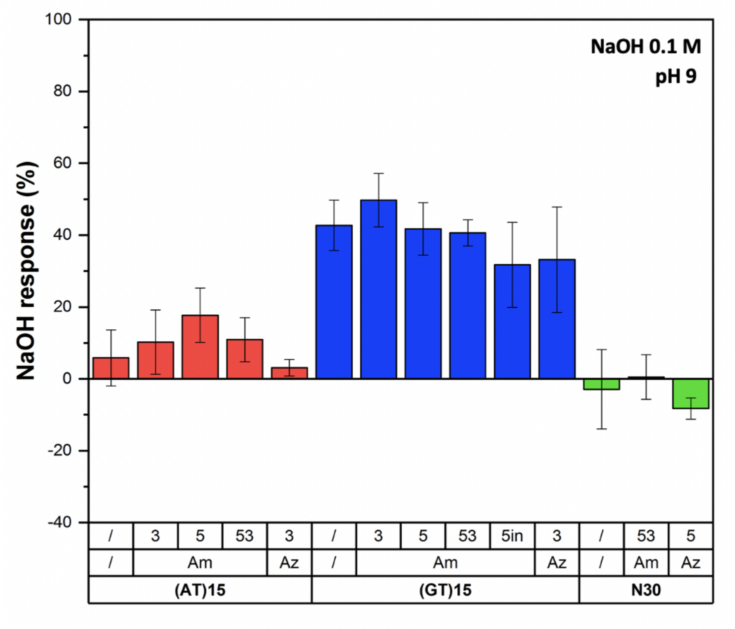


Figure 35: Intensity response following the addition of NaOH 0.1 M to increase the pH of the solution to pH 9. Comparison of the response for modified and unmodified DNA-SWCNTs prepared using MeOH assisted surfactant exchange. Intensity changes were calculated as $I_f$-$I_0/I_0$ for the (6,5) chirality peak (final pH: 9, excitation: 575 nm, incubation period: 10 min). Error bars represent $1\sigma$ standard deviation (n = 3 technical replicates).

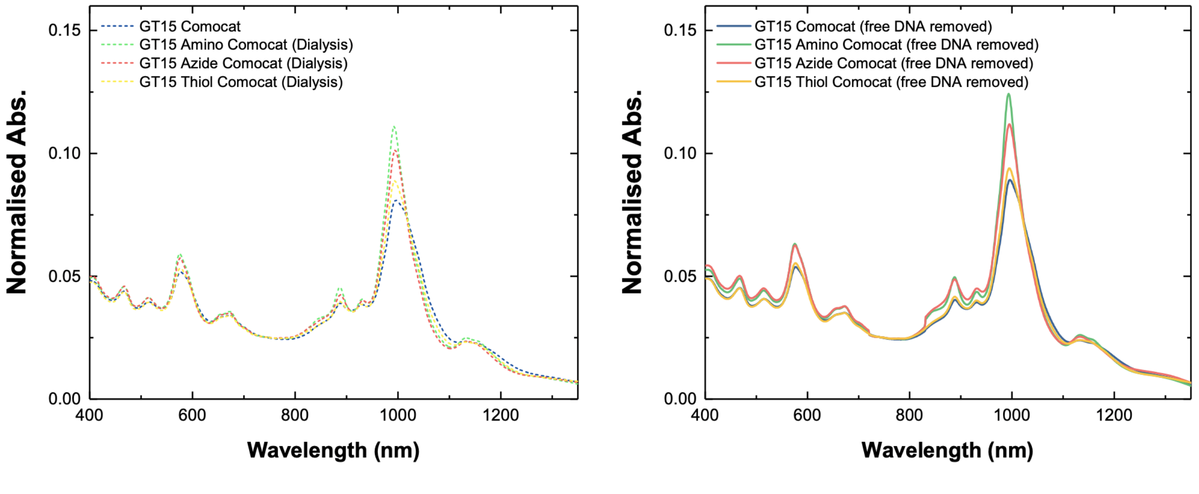


Figure 36: Difference between absorbance in the presence of free DNA and in the absence of free (unbound) DNA. Removing free DNA increased the relative absorbance for each sequence and decreased the FWHM of the peaks indicating improved suspension quality.

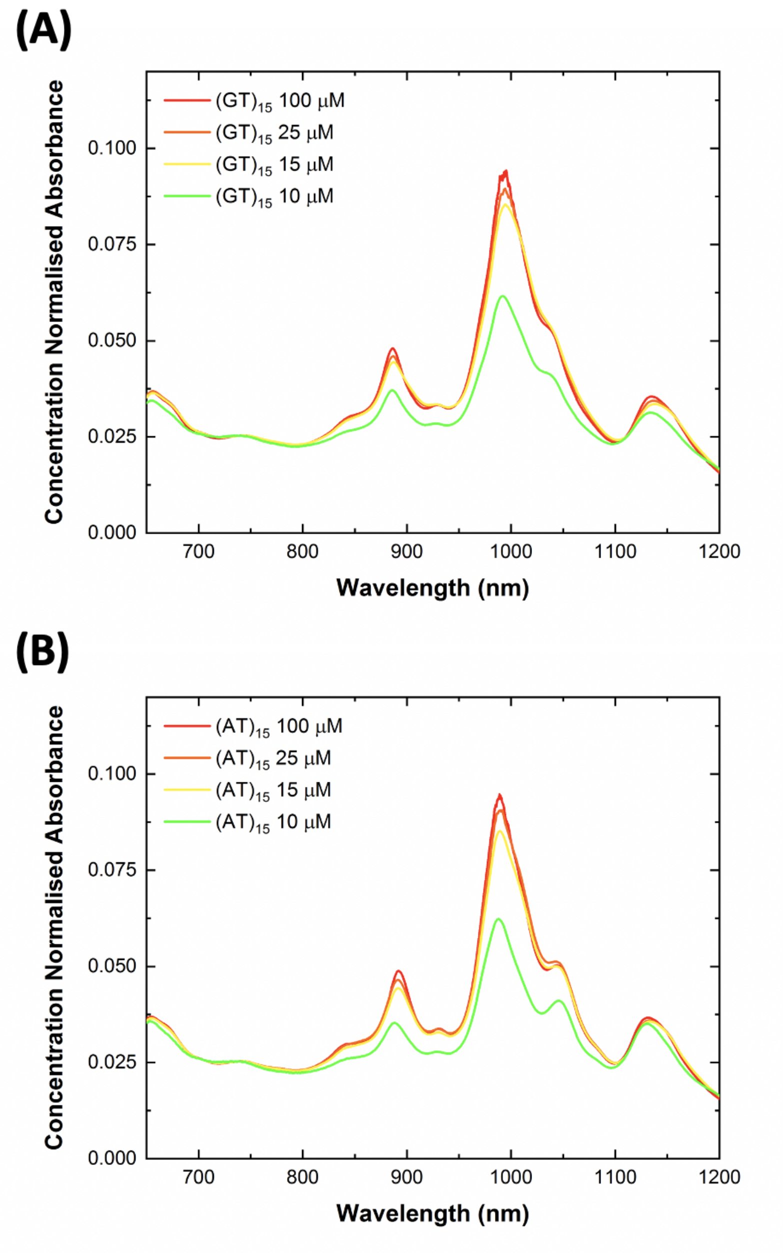


Figure 37: Effect of starting DNA concentration on the dispersion quality for MeOH-assisted surfactant exchanged SWCNT suspensions. The surfactant exchange protocol was carried out on the same batch of SC-SWCNTs using four different concentrations of both **(A)** $(GT)_{15}$ and **(B)** $(AT)_{15}$ to give final DNA concentrations of 100 $\mu$M, 25 $\mu$M, 15 $\mu$M, and 10 $\mu$M. Although a significant decrease in the dispersion yield and quality was observed for both suspensions prepared using 10 $\mu$M final DNA concentration, no significant increases were observed as the concentration was increased above 15 $\mu$M final concentration. For this reason, 15 $\mu$M was selected as the final concentration of DNA used in the MeOH surfactant exchange protocol for this study as it enabled good dispersion of the SWCNTs while reducing any waste of DNA by minimising the excess that remains unbound in the solution. Concentrations below 10 $\mu$M were also tested (7.5 $\mu$M and 5 $\mu$M) however no stable DNA-SWCNT solutions were obtained.

**Table 5:** Estimated relative increase in penetration depths for modified $(GT)_{15}$-SWCNTs compared to unmodified $(GT)_{15}$-SWCNTs. All samples were prepared via direct sonication. Calculations were based on the findings of Lee *et al.*[3] using the equations presented by Lambert *et al.*[4] and an extinction coefficient, $\gamma_{\text{em}}$, of 10 $cm^{-1}$. The increase in penetration depths are shown for the (6,5) (excitation: 575 nm), (7,5) (excitation: 660 nm) and the integrated fluorescence intensity under the 575 nm excitation.

| **(6,5)** | | |
|---|---|---|
| **DNA Sequence** | $\alpha$ | **Δ penetration depth** |
| Am3'-$(GT)_{15}$-SWCNT | 1.55 | 191.5 $\mu$m |
| Am5'-$(GT)_{15}$-SWCNT | 1.80 | 254.4 $\mu$m |
| Am5'3'-$(GT)_{15}$-SWCNT | 1.29 | 109.6 $\mu$m |
| Am5'Int-$(GT)_{15}$-SWCNT | 1.59 | 200.6 $\mu$m |
| Am5'3'Int-$(GT)_{15}$-SWCNT | 3.07 | 487.4 $\mu$m |
| Azide3'-$(GT)_{15}$-SWCNT | 1.68 | 225.1 $\mu$m |
| Azide5'-$(GT)_{15}$-SWCNT | 1.87 | 270.7 $\mu$m |

| **(7,5)** | | |
|---|---|---|
| **DNA Sequence** | $\alpha$ | **Δ penetration depth** |
| Am3'-$(GT)_{15}$-SWCNT | 2.37 | 373.9 $\mu$m |
| Am5'-$(GT)_{15}$-SWCNT | 2.32 | 365.7 $\mu$m |
| Am5'3'-$(GT)_{15}$-SWCNT | 1.91 | 280.2 $\mu$m |
| Am5'Int-$(GT)_{15}$-SWCNT | 1.89 | 276.6 $\mu$m |
| Am5'3'Int-$(GT)_{15}$-SWCNT | 3.53 | 547.8 $\mu$m |
| Azide3'-$(GT)_{15}$-SWCNT | 1.77 | 248.8 $\mu$m |
| Azide5'-$(GT)_{15}$-SWCNT | 2.28 | 357.1 $\mu$m |

| **Integrated Intensity (ex. 575nm)** | | |
|---|---|---|
| **DNA Sequence** | $\alpha$ | **Δ penetration depth** |
| Am3'-$(GT)_{15}$-SWCNT | 1.71 | 232.2 $\mu$m |
| Am5'-$(GT)_{15}$-SWCNT | 1.87 | 272.2 $\mu$m |
| Am5'3'-$(GT)_{15}$-SWCNT | 1.44 | 158.6 $\mu$m |
| Am5'Int-$(GT)_{15}$-SWCNT | 1.62 | 209.5 $\mu$m |
| Am5'3'Int-$(GT)_{15}$-SWCNT | 2.72 | 434.1 $\mu$m |
| Azide3'-$(GT)_{15}$-SWCNT | 1.64 | 213.8 $\mu$m |
| Azide5'-$(GT)_{15}$-SWCNT | 1.88 | 274.8 $\mu$m |
